# Field testing of an enzymatic quorum quencher coating additive to reduce biocorrosion of steel

**DOI:** 10.1101/2022.12.02.518914

**Authors:** Siqian Huang, Celine Bergonzi, Sherry Smith, Randall E. Hicks, Mikael H. Elias

## Abstract

Microbial colonization can be detrimental to the integrity of metal surfaces and lead to microbiologically influenced corrosion (MIC). Biocorrosion is a serious problem for aquatic and marine industries in the world and severely affects the maritime transportation industry by destroying port infrastructure and increasing fuel usage and the time and cost required for maintenance of transport vessels. Here, we evaluate the potential of a stable quorum quenching lactonase enzyme to reduce biocorrosion in the field. Over the course of 21 months, steel samples coated with lactonase-containing acrylic paint were submerged at two different sites and depths in the Duluth Superior Harbor (DSH; Lake Superior, MN, USA) and benchmarked against controls, including the biological biocide surfactin. In this experiment, the lactonase treatment outperformed the surfactin biocide treatment and significantly reduced the number of corrosion tubercles (37% (p<0.01)) and the corroded surface area (39% (p<0.01)) as compared to the acrylic-coated control coupons. In an attempt to evaluate the effects of signal disruption of surface microbial communities and the reasons for lower corrosion levels, 16S rRNA sequencing was performed and community populations were analyzed. Interestingly, surface communities were similar between all treatments, and only minor changes could be observed. Among these changes, several groups including sulfate-reducing bacteria (SRBs) appeared to correlate with corrosion levels, and more specifically, SRBs abundance levels were lower on lactonase treated steel coupons. We surmise that these minute community changes may have large impacts on corrosion rates. Overall, these results highlight the potential use of stable quorum quenching lactonases as an eco-friendly antifouling coating additive.

## Introduction

Biocorrosion is a major issue for the maritime industry all around the world, and it accounts for more than 20% of all corrosion, with an estimated direct cost of $30 to $50 billion each year. [1–3]. Biocorrosion or microbiologically influenced corrosion (MIC) is caused by the metabolic reaction products of microorganisms that are capable of colonizing the metal or metal alloy surfaces. The colonization process results in the formation of a firmly adhering, complex microbial community known as a biofilm. These communities, which can induce biofouling, are harmful to their substrates [4,5] and cause metal surface biodeterioration.

Biocorrosion has a significant impact on steel structures in the Duluth-Superior Harbor (DSH, Minnesota, USA), where this study was undertaken. In the DSH, located on Lake Superior, approximately 20 kilometers of steel sheet piling may be affected by biocorrosion [6]. Significant steel infrastructure corrosion in the DSH was documented, and the steel loss of the harbor due to the extreme corrosion is 2 to 12 times higher than in a comparable freshwater environment [6–8]. These high corrosion rates were related to accelerating biocorrosion rates [6,8,9]. High biocorrosion levels were previously associated with surface colonizing bacteria such as sulfate-reducing bacteria (SRB) and iron-oxidizing bacteria (IOB) [9–11]. Like many other freshwater and marine environments, corroded steel surfaces and tubercles in the DSH are covered by complex microbial biofilms containing bacteria responsible for the corrosion of steel [8,12–17]. Within these surface biofilms, SRB live in the anoxic zone of the corrosion tubercles and can accelerate corrosion by either producing hydrogen sulfide that reacts with iron forming FeS or using iron directly in the metabolism as an electron donor [13].

Solutions used to combat biocorrosion predominantly rely on biocides [18–21]. While these biocidal compounds can delay biofilm formation, they are harmful to the environment and can accumulate. Recent efforts to address the ecological and economic concerns associated with biocorrosion have focused on biological or benign molecules to combat biocorrosion [20–22]. Molecules preventing bacterial adhesion or biofilm formation were tested in different coating chemistries and on different substrates, including metals [18,21,23], silicon, organic polymers, and glass [21]. The tested molecules include an underexplored class of biological molecules, namely enzymes. While most enzymes lack the necessary stability to retain activity in coatings, some, particularly those encoded in extremophiles, may remain active as a coating additive. Specifically, we recently investigated the ability of quorum quenching lactonases embedded in materials such as coatings to remain active under harsh conditions [24,25]. These lactonases are capable of interfering with Acyl Homoserine Lactone (AHL) based microbial communication by degrading these molecules [26–28]. AHL signaling is used by numerous microbes to regulate behaviors such as biofilm formation in response to cell density, a communication system called quorum sensing (QS) [29]. By disrupting QS, lactonase enzymes can inhibit bacterial virulence and the formation of bacterial biofilms in the context of biofouling [13,30,31]. Interference in QS can reduce microbiologically influenced corrosion, possibly by reducing the amount of surface biofilm [32].

We previously evaluated the potential of various compounds, including a quorum quenching lactonase, to inhibit biocorrosion in a laboratory setting [33]. These studies were made possible by the identification [34,35], characterization [24] and engineering of exceptionally stable lactonase enzymes [36,37]. Surprisingly, the lactonase coating additive showed greater reduction of corrosion than other tested biocides such as surfactin and magnesium peroxide. This finding is intriguing because, in contrast to the biocidal molecules that dominate the antifouling and anticorrosion coatings market [18,19], these stable lactonase enzymes show low to no toxicity [38,39], and little to no effect on bacterial fitness [24,40]. Lactonases do not need to enter microbial cells, and do not need to bind to a receptor [41,42]. Instead, they act by hydrolyzing signaling molecules secreted into the medium, reducing the concentration of the signal and thereby affecting bacterial behavior and inhibiting biofilms [27,28,43], including the associated complex microbial communities [39]. Intriguingly, the inhibition of the corrosion observed for the lactonase additive was concomitant to significant changes in the bacterial community composition [44]. This observation is consistent with other studies highlighting the ability of AHL signal disruption to alter microbial community structures [25,45,46].

We report the evaluation of the ability of a lactonase enzyme used as an additive in an acrylic coating to reduce biocorrosion, and compared it with a coating containing a biocidal additive (surfactin) in a field experiment. Quantification of corrosion at different sampling times showed that the use of lactonase as a coating additive lead to significantly higher inhibition of biocorrosion over the project period (21 months). Analysis of the surface microbial population structure revealed that the presence of the lactonase induces changes to the surface community, specifically for bacterial orders of *Rhizobiales, Holophagales, Sulfuricellales, Desulfovibrionales*, and *Desulfuromonadales*. These results suggest a quorum quenching lactonase used as a coating additive can be a reasonable means to reduce biocorrosion, including over a significant period of time (21-months).

## Materials and Methods

### Experimental Coupon Construction and Pretreatment

Steel coupons (11.4×4.8×0.95 cm) were cut from hot rolled ASTM-A328 steel, the same material used to build the steel sheet pilings that support most of the docks and bulkheads in the Duluth-Superior Harbor (DSH) and many other ports. Before treatment, the coupons were washed with soapy water, lightly brushed, and rinsed with Milli-Q water. Each coupon was given a unique number, weighed, wrapped in aluminum foil, and autoclaved before being assigned to a specific experimental treatment at random.

### Coupon Treatments and Installation

Two treatment sets and two control sets of the steel coupons were investigated for corrosion rate and bacterial communities in this field experiment. They were installed in the DSH by hard hat divers on Aug 22, 2017 and were planned to be exposed in the harbor environment for a period of 1, 2, 8 and 21 months. The treatments are: A, bare steel control without coating; B, acrylic coating control; C, acrylic coating with 200 μg/mL surfactin; D, acrylic coating with 200 μg/mL lactonase enzyme.

For each treatment (A, B, C, D) at each test site and submerged depth, duplicate coupons were installed into electrically isolated trays with all the scratched surfaces facing the same side (**Fig. 1A**). Metal tray frames that contained the sheet steel coupons (**Fig. 1B**) were attached at 1 m and 3 m below the waterline to corroding structures in two sites (Hallett Dock 5 and LaFarge Dock) in the DSH. Each frame held eight of the coupon trays. A total of 128 coupons in 32 trays were installed.

**Fig 1.**
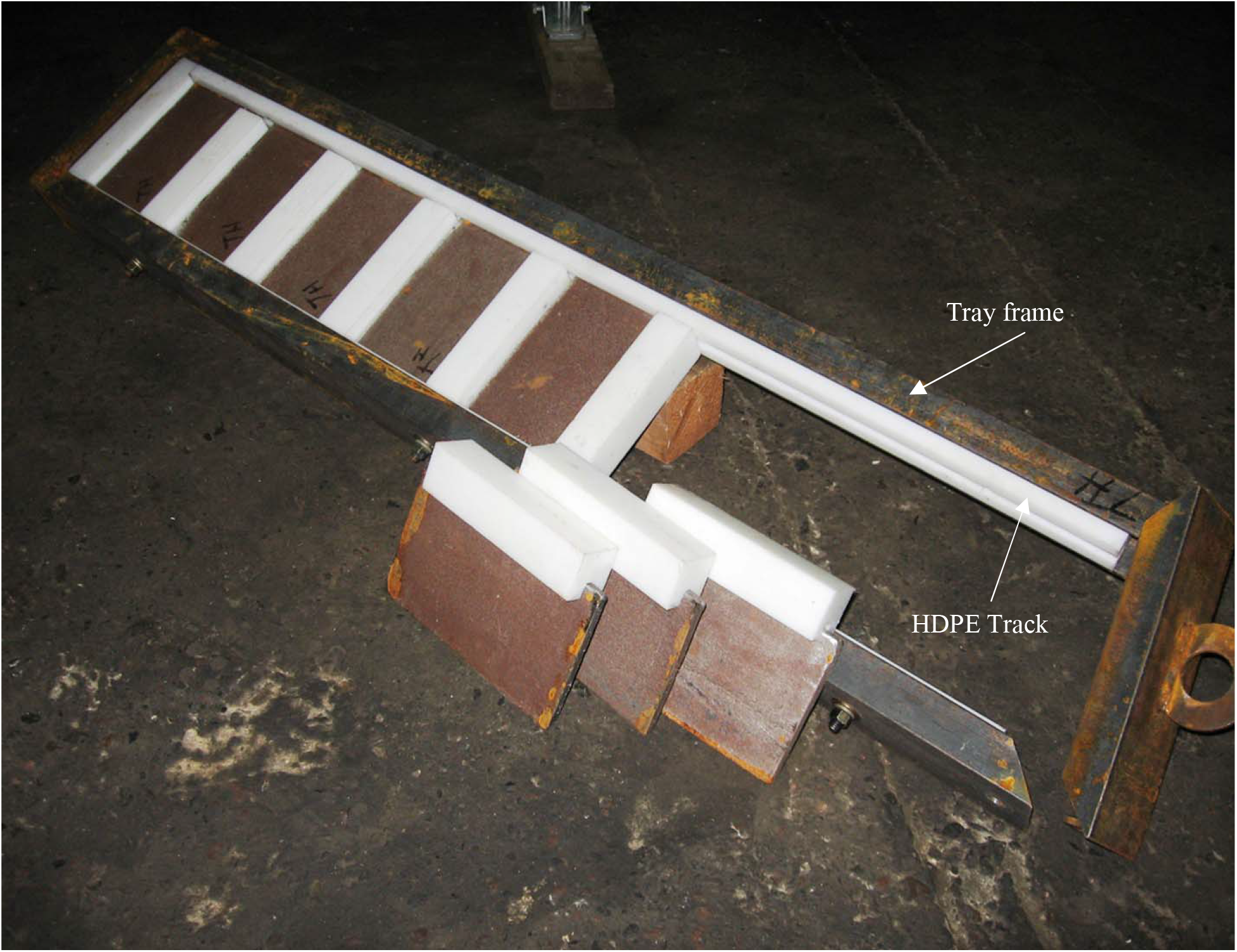
Steel coupon installation. The coupon trays were installed in the frame by inserting the coupons into the HDPE tracks and sliding to their positions.

### Production and purification of the lactonase enzyme

The SsoPox W263I was produced as previously described [33,44]. Briefly, *E. coli* strain BL21(DE3)-pGro7/GroEL (Takara Bio) was grown in 500 mL of ZYP medium [47] (supplemented with 100 μg/mL ampicillin, 34 μg/mL chloramphenicol) and 0.2% (w/v) arabinose (Sigma-Aldrich) was added to induce the expression of the chaperones GroEL/ES [44]. Lactonase purification was carried out as previously described [44]. The high thermal stability of the enzyme allows for its purification using a 30 min heating step at 70 °C, followed by differential ammonium sulfate precipitation, dialysis, and exclusion size chromatography. Pure *Sso*Pox W263I enzyme samples were quantified using a spectrophotometer (Synergy HTX, BioTek, USA) at OD_280nm_ and a protein molar extinction coefficient as calculated by PROT-PARAM (Expasy Tool software) [48].

### Coupon coating treatments with coating scratch

Coating treatments were prepared using commercial Sher-Clear 1K - Acrylic Clear Coat (Sherwin-Williams, OH, USA). Six mg of surfactin (Sigma Aldrich Corp. St. Louis, MO, USA) or 6 mg of lactonase enzyme were added to 30 ml of acrylic base to create 0.2 mg/mL coating treatments. We note that while the concentration for surfactin and lactonase are identical, they are not equimolar due to the large molecular weight difference between the molecules. Indeed, the molecular weight of surfactin is 1,036.3 g/mol, whereas it is 35,491.89 g/mol for the lactonase *Sso*Pox W263I, or a ∼34-fold molar excess of surfactin relative to the lactonase.

The steel coupons were dipped into the prepared coatings for 1 min then air-dried at room temperature overnight. Each coupon was covered in a uniform layer of acrylic coating on all surfaces. All coating-treated coupons were scratched on one side with a steel file. Each scratched surface had two parallel scratches from side to side (**Fig. S1**). The width of all scratches was approximately 3 mm.

### Coupon collection and tubercle analysis

Experimental and control coupons were retrieved after approximately 1, 2, 8 and 21 months of exposure (September 2017, October 2017, May 2018, June 2019) in the DSH. Sample analysis was performed as described in the preceding study [33]. Coupons in each treatment were photographed with a digital camera immediately after being removed from the harbor water. The number and coverage of corrosion tubercles were used to assess the intensity of biocorrosion. Tubercle numbers were manually counted, and the total tubercle area was measured using ImageJ (NIH, Bethesda, MD, USA). The tubercle numbers, tubercle coverage, and surface roughness were analyzed using Microsoft Excel software (one-way, non-paired T-tests between each treatment and control).

### Biological Analysis Methods

DNA extraction was performed as described in the preceding study [33]. Each coupon’s material, including biofilm and tubercles, was scraped into a sterile 50 ml polypropylene centrifuge tube (Corning, New York, USA) using a stainless-steel spatula. DNA was extracted from a 0.5 g subsample of this surface material from each coupon using the PowerSoil DNA kit (MoBio Laboratories). The extracted DNA was used to sequence the V4 region of the 16S rRNA gene and study changes in bacterial community composition. We chose the V4 region of the 16S rRNA gene because the Illumina (San Diego, CA, USA) MiSeq 2×300 16S rRNA V4 protocol yields a greater number of high-quality overlapped pair-end sequence reads than other 16S rRNA gene variable regions (i.e. V1-V3) [49]. Furthermore, even with relatively short reads from the 16S rRNA V4 region, good resolution can be obtained at the order through family taxonomic levels [50]. The extracted DNA was quantified using a NanoDrop 2000 Spectrophotometer (Thermo Scientific, Waltham, MA, USA) before being analyzed in a qPCR assay and partial 16S rRNA gene sequencing at the University of Minnesota Genomics Center.

In addition, quantitative PCR (qPCR) experiments were performed to determine the abundance of sulfate-reducing bacteria (SRB) in all samples using the dissimilatory sulfite reductase (*dsr*A) gene as a probe. SRB abundance was estimated using a modified procedure from Kondo et al. [51] by quantifying copies of the *dsr*A gene. Quantitative PCR was performed in a 25 μL reaction volume consisting of 12.5 μL Brilliant II SYBR Green Master Mix (Agilent Technologies), 1.0 μL of 10 μM forward DSR-1F (5’-ACS CAC TGG AAG CAC GGC GG −3’) and reverse DSR-R (5’-GTG GMR CCG TGC AKR TTG G −3’) primers, 2.0 μL of 10 mg/ml bovine serum albumin, 3.5 μL of nuclease-free water, and 5.0 μL of DNA template (10 ng total) on a Rotor-Gene 3000 (Corbett Life Science, NSW, Australia) qPCR thermal cycler. A standard curve was developed using *Desulfovibrio vulgaris* subsp. vulgaris genomic DNA (ATCC 29579D-5) amplified with the DSR1F and DSR-R primer set. The standard curve ranged from 0.1 pg to 0.1 ng (400 to 4 × 10^8^ copies of the *dsr*A gene) of this genomic DNA. On each sample, the qPCR analyses were carried out in triplicate.

### DNA sequence processing and analysis

DNA sequencing and data analysis was performed as previously described [33]. DNA from microbial communities on the surface of all steel coupons was shipped overnight and sequenced at the University of Minnesota Genomics Center using an Illumina MiSeq, which produced the sequence of the 254 bp portions of the 16S rDNA V4 region. llumina sequence data from this study were submitted to the NCBI BioProject database under accession number PRJNA859427. In each analysis, thirty samples were multiplexed in order to obtain approximately 500,000 sequences per replicate coupon sample. Sequencing data was processed and analyzed using MOTHUR [52]. Sequence reads with ambiguous bases, homopolymers >7 bp, more than one mismatch in the primer sequence, or an average per base quality score of less than 25 were removed. Sequences that appeared only once in the total set were assumed to be the result of a sequencing error and were removed prior to further analysis. Chimeric sequences were also removed using the UCHIME algorithm [53]. The remaining sequences were clustered into operational taxonomic units (OTUs) using a cutoff value of 97 percent. The Ribosomal Database Project (RDP) taxonomic database was used to assign taxonomy to OTU consensus sequences. MOTHUR was also used to compute coverage and generate a Bray-Curtis dissimilarity matrix. Bacterial communities from various samples were compared in MOTHUR using ANOSIM, a nonparametric procedure that tests for significant differences between groups by using Bray-Curtis distance matrices. Bacterial communities on tubercles from different treatments were compared using PC-nonmetric ORD’s multidimensional scaling (NMDS) ordinations (MJM Software Designs, Gleneden Beach, OR USA).

The Linear discriminant analysis (LDA) Effect Size (LEfSe) analysis [54] was used to identify bacterial orders that are responsive to the different treatments in the four sampling periods. To find species with significant variations in abundance across groups, the Kruskal-Wallis (KW) sum-rank test was performed. Then, the rank sum test was performed to assess the differences across groups. Linear discriminant analysis (LDA) was used to reduce the dimension and quantify the differences. All sites and depths are combined in the analysis.

Relative abundance is significant when P < 0.05, logarithmic LDA score ≥2.

## Results and Discussion

### Lactonase as a coating additive can reduce biocorrosion of steel in field studies

We previously investigated the ability of lactonase as a coating additive to reduce biocorrosion in a controlled laboratory environment over 8 weeks of submersion [33]. Formulation of enzymatic coatings was possible because of the exceptional stability of the lactonase enzyme Ssopox W263I. This enzyme was identified from the thermophilic archaeon *Saccharolobus solfataricus* [34,55,56]. The *Sso*Pox enzyme, and particularly its mutant W263I [36], possesses exceptional thermal stability, as well as solvent, protease, and aging resistance [24]. Here, we conducted a follow-up study by challenging an enzymatic coating in the field, over nearly two years.

Sample coupons were retrieved after 1, 2, 8, and 21 months of submersion in the Duluth-Superior harbor. Visible corrosion occurred on the steel coupons within 1 month of submersion in the DSH water. Photographs showed clear signs that corrosion tubercles started forming and were growing on the uncoated control and the scratched area of the steel surfaces in the early stage of the experiment (**Fig 2**). By the end of the field experiment (21 months), both the uncoated control and acrylic-only control coupons were heavily covered by tubercles (**Fig 2; Fig. S2**). Significant biofouling, including mussels, can also be noted on the acrylic coating control samples.

**Fig 2.**
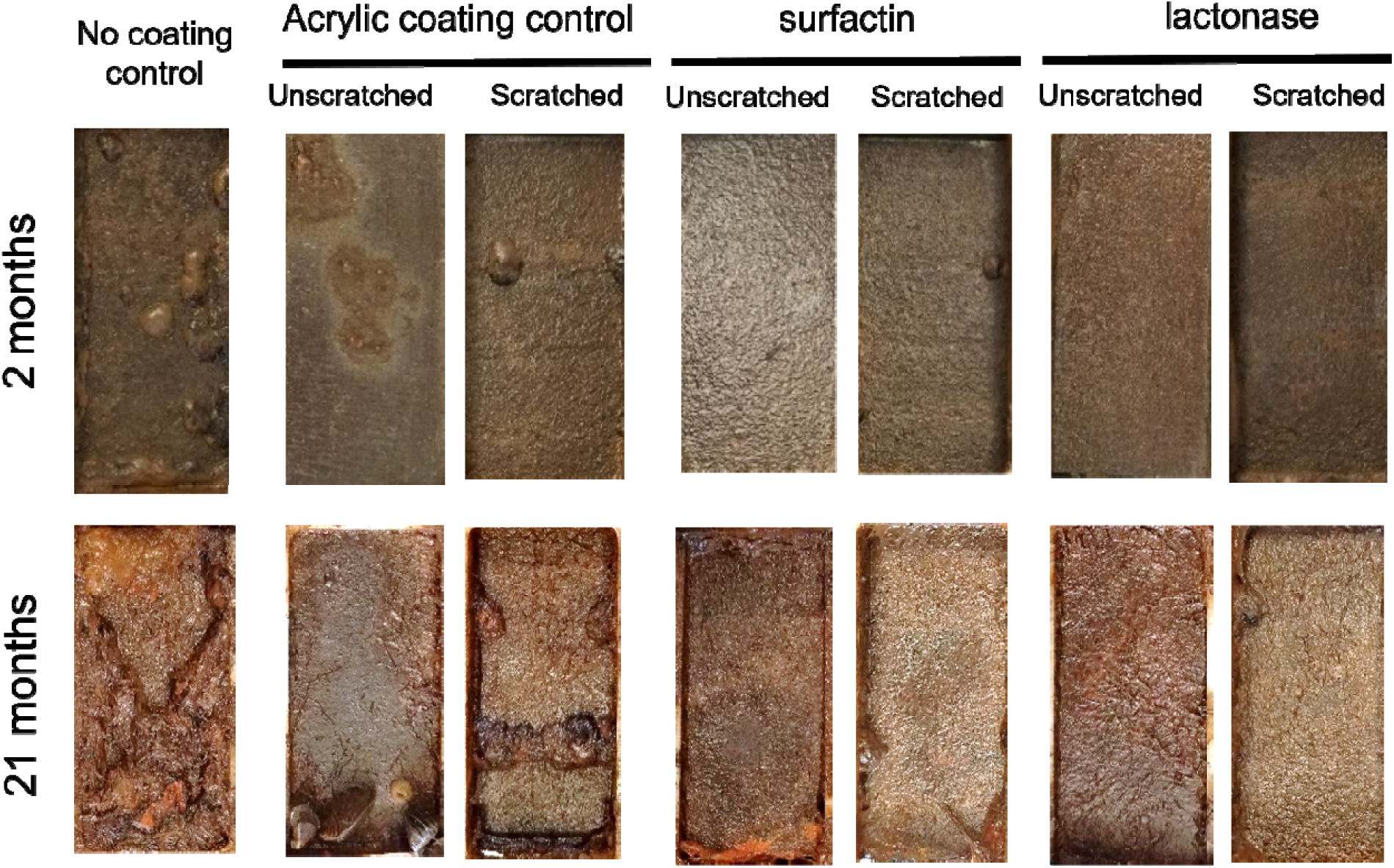
Photographs of control and treated steel coupons after 2 (upper) and 21 (lower) months of exposure in the Duluth-Superior Harbor (Site: HD5, Depth: 3m). Images of the coupons after 1 and 8 months are included in **Fig. S2**. Images on the right in each test group show the scratched face of the sample coupon, with two parallel scratches (see methods and **Fig. S1** for description)

Sample corrosion was examined by counting corrosion tubercule numbers (**Fig. 3**) and determining the fraction of the corroded surface area (**Fig. 4**) on the steel coupon surfaces. Corrosion rates were slightly higher for coupons submerged at 3 m depth compared to coupons incubated at 1 m depth. Moreover, corrosion rates were similar between the two sampling sites in the harbor; Hallett Dock 5 and LaFarge Dock (**Fig. S3&S4**). As expected, the bare steel control showed the highest corrosion, *i*.*e*. highest numbers of tubercles and corroded surface, with an average of 29 tubercles and 63% of the surface area covered with tubercles after 21 months (**Fig. S3**). As expected, scratched surfaces showed higher corrosion than unscratched areas, confirming that scratching coating surfaces accelerated the corrosion process (**Fig. 3&4**).

**Fig 3.**
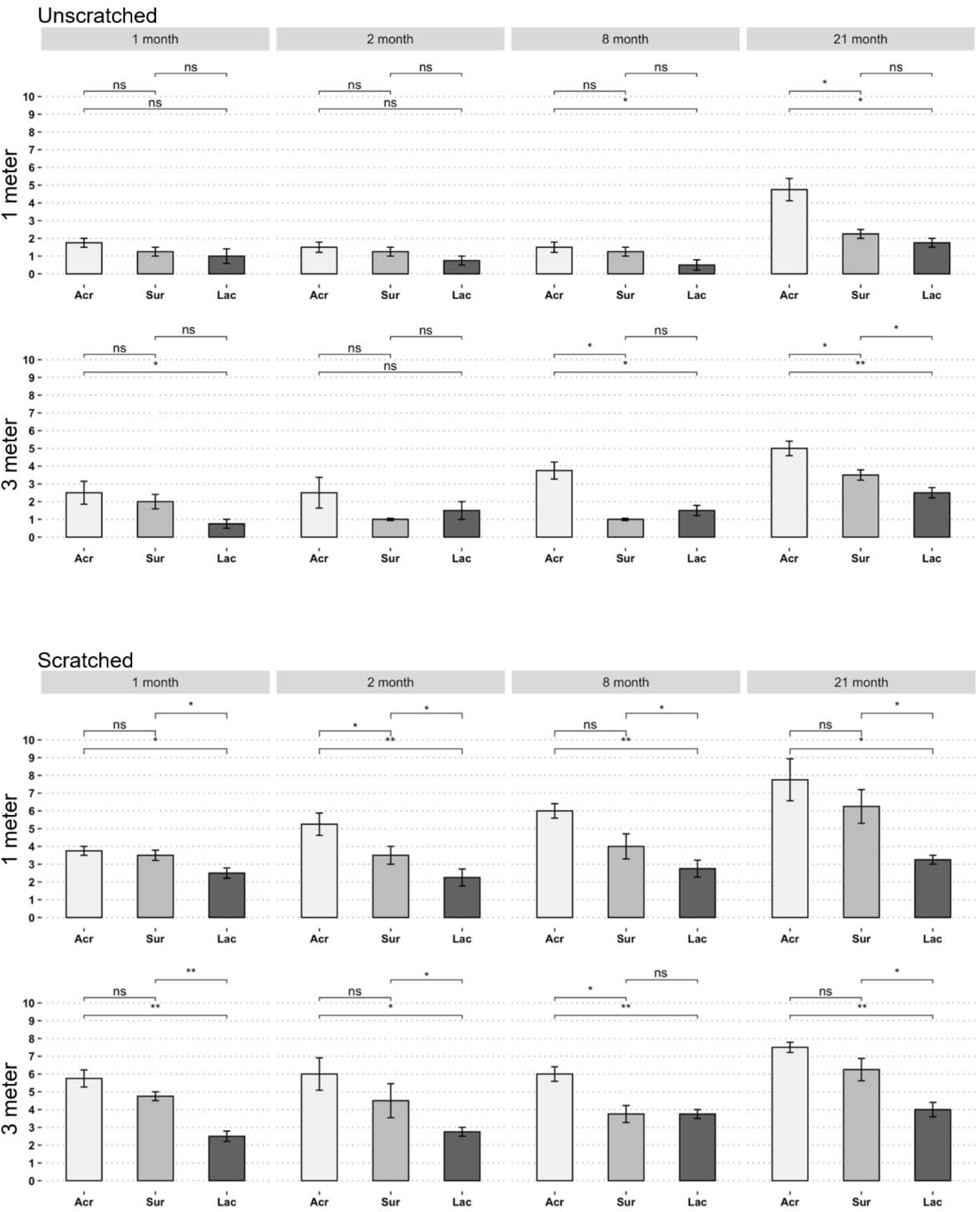
Corrosion tubercule counts revealed a reduction of corrosion with lactonas treatment. Counts of corrosion tubercles on unscratched (upper panel) and scratched (lower panel) steel coupons with different experimental treatments. Mean values for 2 coupons at both sites are shown (n=4). The tested conditions are acrylic coating control (Acr); acrylic coating with 200 ug/ml surfactin (Sur), acrylic coating with 200 ug/ml *Sso*Pox lactonase enzyme (Lac). Means with one or two asterisks were different at the significance levels of p<0.05 or p<0.01, respectively.

**Fig 4.**
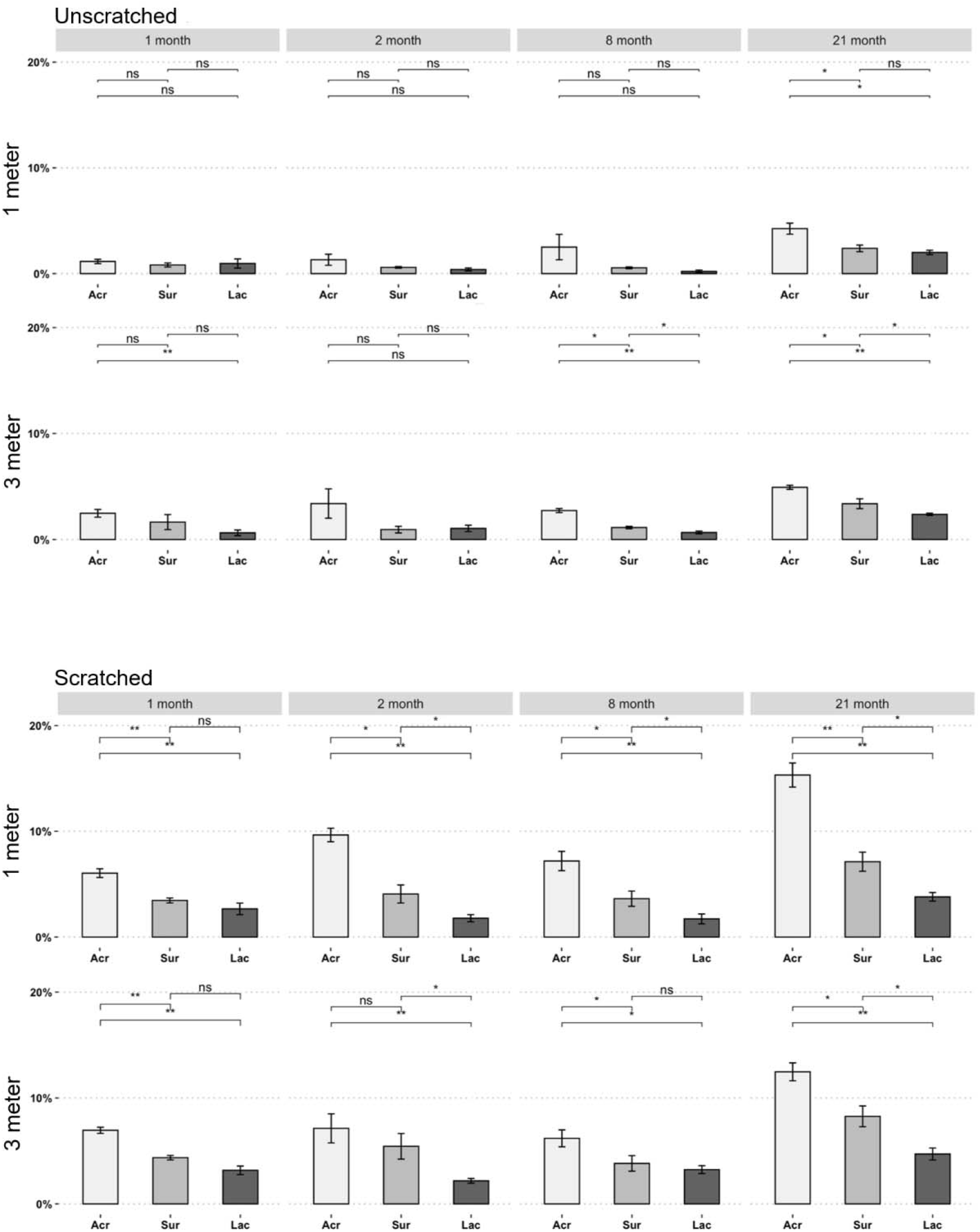
Corrosion tubercule surface coverage revealed a reduction of corrosion with lactonase treatment. Coverage of corrosion tubercles on unscratched (upper panel) and scratched (lower panel) steel coupons with different experimental treatments. Mean values for 2 coupons at both sites are shown (n=4). The tested conditions are acrylic coating control (Acr); acrylic coating with 200 ug/ml surfactin (Sur), acrylic coating with 200 ug/ml *Sso*Pox lactonase enzyme (Lac). Means with one or two asterisks were different at the significance levels of p<0.05 or p<0.01, respectively.

The surfactin treatment, used in this study as the biocide positive control, shows expected reduction of corrosion as previously reported [57]. Both tubercle number and corroded surface area appeared reduced as compared to the no additive control acrylic coating (**Fig. 3&4**). Corrosion tubercle counts with surfactin as an additive was significantly reduced compared to the acrylic coating alone treatment in samples collected after 21 months, unscratched (e.g. reduction by 30% (p<0.05) for samples at 3 m). Scratched samples did not show a significant difference in tubercle count (**Fig. 3)**. Similarly, corroded surface area was significantly reduced with surfactin as an additive after 21 months for both scratched and unscratched surfaces (e.g. reduction by 33% (p<0.05) for samples at 3 m**; Fig. 4**).

Interestingly, the lactonase treatment showed remarkable corrosion inhibition. Both tubercle numbers and corroded surface area were reduced compared with the no additive acrylic coating control for all tested conditions (**Fig. 3&4**). For unscratched samples, reduction of tubercle numbers was significant for month 1 sampling (3 m) and months 8 and 21 sampling times (1 and 3 m), while corroded surface area was significantly reduced for month 1 sampling (3 m), month 8 sampling time (3 m) and month 21 sampling time (1 and 3 m). After 21 months, the reduction of tubercles and corroded surface area are 56% (p<0.01) and 52% (p<0.01), respectively. For scratched samples, all recovered samples at 1 month (1 m) showed significantly reduced tubercle counts and corroded surface areas. For scratched surfaces, after 21 months, the reduction of tubercles and corroded surface area are 52% (p<0.01) and 69% (p<0.01), respectively.

Compared to the surfactin treatment, the lactonase coating also showed greater corrosion inhibition over the course of the experiment. The lactonase treatment showed significant reductions in corrosion tubercles and corroded surface compared to the surfactin treatment for both 8- and 21-months sampling times (3 m; unscratched). For scratched samples, reductions were even more statistically significant: nearly all sampled conditions showed greater corrosion protection for the lactonase treatment, including sampling times 2, 8 and 21 months (1 and 3 m). For example, in the case of scratched surfaces incubated at 3 m depth and collected after 21 months, the reduction of tubercles and corroded surface area for the lactonase treatment compared to the surfactin treatment were 36% (p<0.05) and 43% (p<0.05), respectively. The observed reduction of microbiologically induced corrosion is consistent with previous reports [32]. In fact, some key SRBs are known to produce AHLs, and the latter can stimulate sulfide oxidation by SOBs [20]. A lactonase containing coating, therefore, would be expected to disrupt these mechanisms.

Over the course of this experiment, sample coupons also accumulated a limited amount of macrofouling (**Fig. 2, S5-S8**). Unfortunately, possibly due to the stochastic nature of mussel attachment, the differences between treatments were not statistically significant, even if results appear to indicate a reduction of the attachment of mussels to the substrate in the lactonase treatment, as compared to other tested coatings. Consistently, total mussel weight was also lower for the lactonase treatment (**Fig. S6&S8)**. Further studies will be needed to confirm this initial observation.

The results described here are consistent with our seminal observations on biocorrosion [32], but also with other reports [32]. While AHL-based signaling is used by numerous bacteria and AHL signals are sensed by a wide bacterial audience [58], approaches pertaining to AHL signal disruption may have long been hypothesized to only affect AHL-producing / sensing bacteria. More recent work on very different biofilms (e.g. water treatment [46] or soil community [25] are consistent with observations in the current study. These observations that AHL signal disruption inhibits the formation of complex biofilms suggest that this molecular communication system also has profound effects on microbial communities and their phenotypes. It is unclear however whether these community-wide effects are due to the direct change in the behavior of bacteria sensing AHLs, the latter behavioral change causing indirect modulation of the rest of the microbial population. These effects could also be explained by a larger than anticipated fraction of microbes in communities responding to AHLs, causing signal disruption to directly alter their behavior. Or it could be a combination of direct effects on bacteria using AHLs for signaling, indirect effects on other bacteria whose behavior adapts to the change in AHL-using bacteria, and a larger, uncharacterized portion of bacteria capable of sensing and responding to these changes.

### Alteration of surface microbial community by lactonase-containing coating

In order to better understand the effects of AHL signal disruption on the surface of the treated steel, we undertook Illumina 16S rDNA sequencing and analyzed surface microbial community composition. When comparing all sequenced samples, this analysis indicated that most of the observed divergence in surface microbial population was caused by sampling at different sites (AMOVA =2.42668, p = 0.02), different depths (AMOVA = 5.5809, p = <0.001) and different sampling times (p-values all <0.007; **Table S1; Fig. 5, 6, S9, S10**). These differences were unrelated to the treatments and are therefore not the focus of the analysis below. However, they highlight the necessity to evaluate nonbiocidal antifouling molecules at a range of sites, in order to account for any differences in local microbial ecology, and validate additive-containing coatings as an antifouling technology that can operate in different microbial environments.

**Fig 5.**
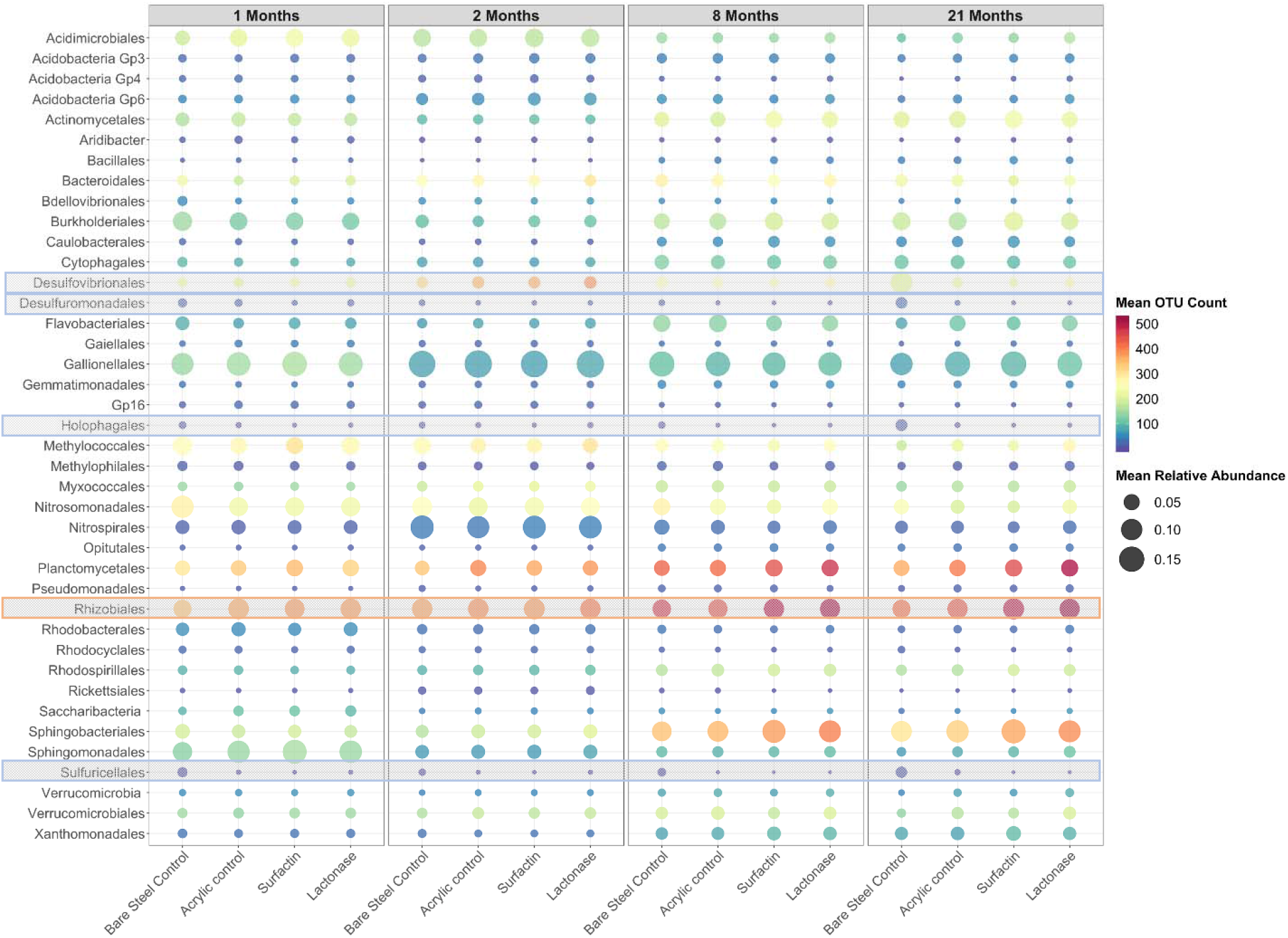
Microbial surface community analysis. Heatmap comparing the relative abundances of partial 16S rRNA sequences and OTU richness for the top 40 bacterial orders in all samples from each treatment and control. Diversity is indicated by the number of OTUs in each bacterial order. The mean was calculated by averaging different site and depth replicate samples (n=8). The orders with the highest relative abundance change are highlighted in boxes. The full heatmap with data separated by time, depth and site is shown in Fig S9.

**Fig 6.**
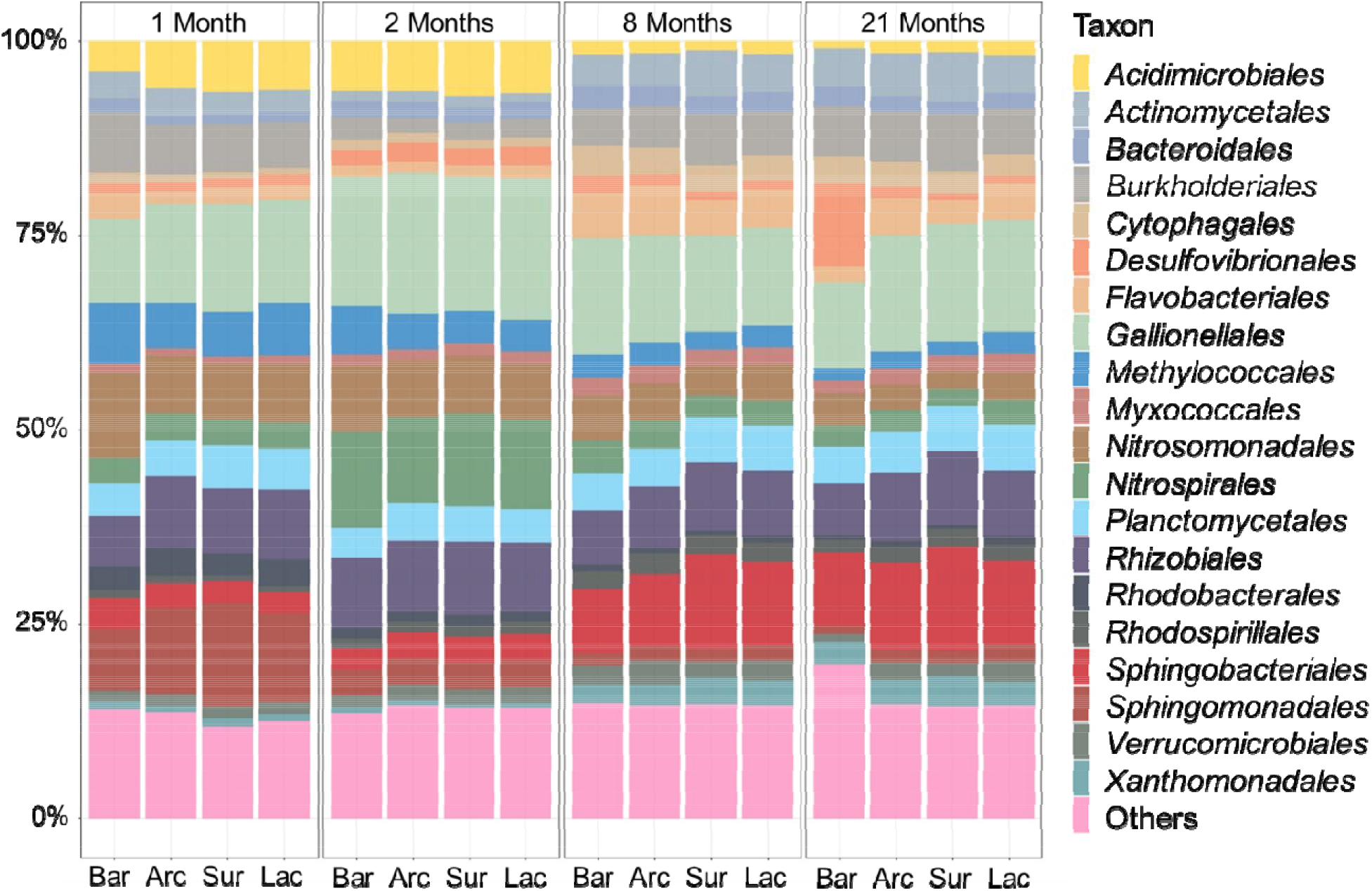
Most common bacterial orders in surface communities. Relative abundance of partial 16S rRNA sequences for the top 20 bacterial orders in all samples from each treatment and control. The mean was calculated by averaging different site and depth replicate samples (n=8). Bar: Bare steel control without coating, Acr: Acrylic coating control, Sur: Acrylic coating with 200 ug/ml surfactin, Lac: Acrylic coating with 200 ug/ml SsoPox lactonase enzyme. The full bar graphs with data separated by time, depth, and site are shown in Fig S10.

In regards to treatments, surface microbial population analysis revealed that the microbial community on the bare steel control was significantly different from any other treatment (p values all <0.001; **Table S1; Fig. S9-S13**). When comparing the surfactin and lactonase treatments to the acrylic base control (considering all samples together) the groups do not show different communities (acrylic control and lactonase: p = 0.213; acrylic control and surfactin: p = 0.409; lactonase and surfactin: p = 0.467; **Table S1**). Differences in bacterial biofilm communities on treated and acrylic control coupons only appeared different after 21 months of submersion (acrylic-lactonase; p = 0.066 (HD5, 3m); p = 0.074 (LAF, 3m), acrylic-surfactin; p = 0.055 (LAF, 1m); p = 0.096 (LAF, 3m)). (**Table S1**). Similarly, the two treatments, lactonase and surfactin, only showed significant surface population differences after 21 months: p = 0.053 (LAF, 1m); p = 0.084 (HD5, 3m) (**Table S1**). Members of the *Gallionellales (14*.*0%), Sphingobacteriales (6*.*3%), Rhizobiales (5*.*6%), Nitrospirales (5*.*4%), and Burkholderiales (5*.*2%)* were the dominant bacterial orders found on all coupons (**Fig. 5, 6, S10**). Between treatments, some orders show noticeably different abundance or OTUs counts, such as *Rhizobiales, Holophagales, Sulfuricellales, Desulfovibrionales*, and *Desulfuromonadales*. These results suggest that despite large differences in corrosion rates, the nature of the surface bacterial community is not dramatically different between treatments, and a large component of the community is shared between treatments. Yet, smaller changes in the abundance of key microbes might be involved in the observed different corrosion rates.

We then attempted to examine the potential relationships between the relative abundances of known iron-oxidizing bacteria and sulfate-reducing bacteria in different treatments. Because treatments showed different corrosion levels (bare steel being the most corroded, lactonase being the least), we could use the different treatments as a proxy for corrosion levels. Intriguingly, when attempting to correlate these bacterial groups to corrosion levels (**Fig. S14&S15**), we found that the relative abundance of *Holophagales, Sulfuricellales, Desulfovibrionales*, and *Desulfuromonadales* correlated with corrosion levels (**Fig. 7**), while the relative abundance of *Rhizobiales* did not correlate with corrosion levels. A further analysis was performed to verify this finding. A linear discriminant analysis (LDA) Effect Size (LEfSe) analysis was performed to identify the specific bacterial taxa with significantly higher abundance in microbial communities for each treatment (**Fig. S16**). When treatment communities were most distinct (at 21 months), it appeared that *Holophagales, Sulfuricellales, Desulfovibrionales*, and *Desulfuromonadales* were among the bacterial orders most characteristic of the highly corroded bare steel samples, while *Rhizobiales* appeared to be an order more characteristic of surface communities on surfactin- and lactonase-treated coupons. Remarkably, the bacterial orders that correlated with tubercle formation are reported to have AHL quorum sensing capability (**Table S2**), and changes in their relative abundances may depend on AHL signaling. While members of the *Rhizobiales* are known to be able to oxidize iron, which has been reported to accelerate corrosion of iron [57,59,60], we unexplainably observed here that the relative abundance of *Rhizobiales* appeared to not correlate with corrosion levels. However, members in *Holophagales* such as *Geothrix fermentans* are well documented as iron (III) reducing bacteria [61]. *Desulfuromonadales and Desulfovibrionales* are major orders of SRB, have been found in biocorrosion samples [62,63] and were observed to correlate with corrosion levels in this study.

**Fig 7.**
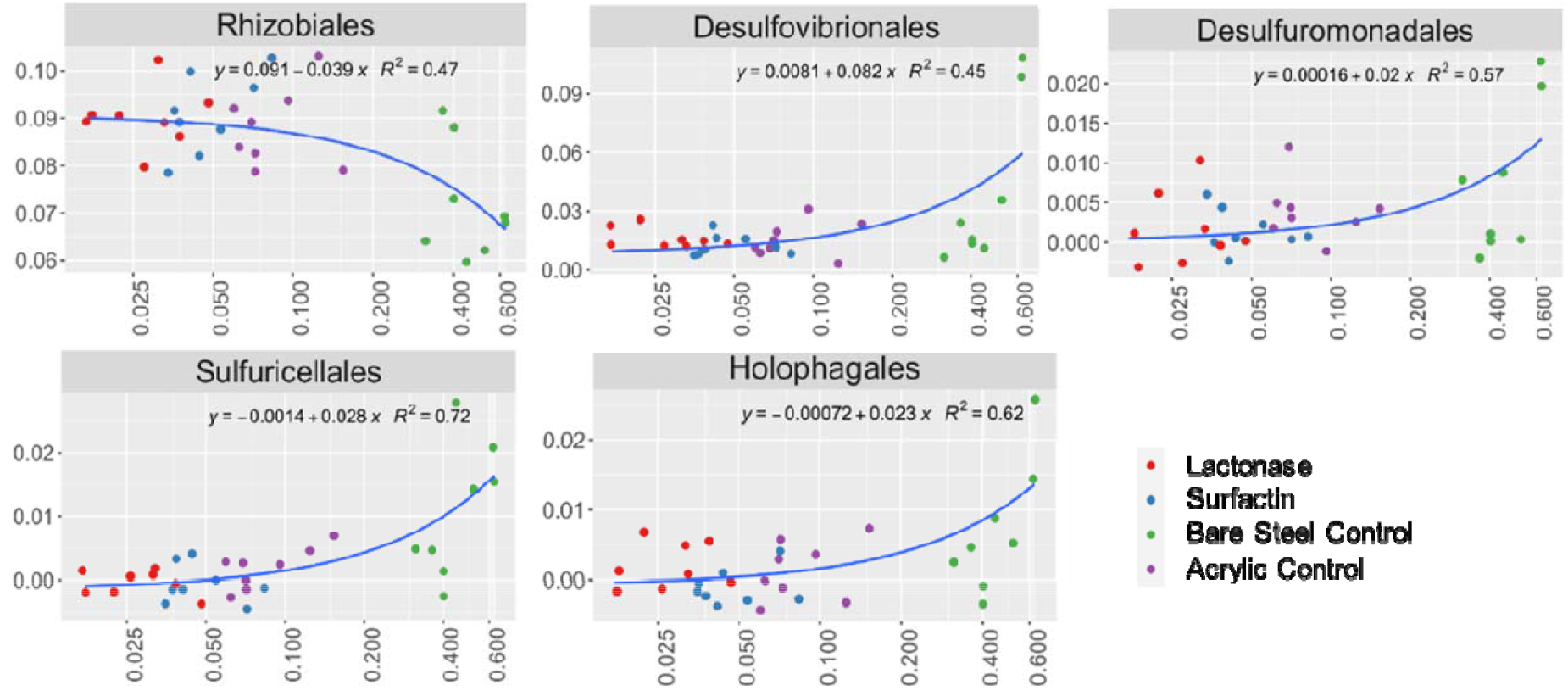
Relative abundance and corrosion levels appear to correlate differently for some SRBs. Log scale plots showing the correlation between relative sequence abundance of 5 bacterial orders and the tubercle coverage for treatment and control samples. Y axis: relative abundance, X axis: tubercle coverage (log scale). The log scale plots of the rest of major orders were shown in Fig S15.

Dissimilatory sulfite reductase (*dsrA*) gene qPCR analysis was performed to complement the 16S rDNA sequence analysis work. This measurement can be used as a proxy to quantify the abundance of SRBs in a sample [51]. The *dsrA* gene quantification performed on samples collected after 8 months of submersion showed reductions of *dsrA* gene abundance on the surfactin and lactonase treated coupons (30% and 46%, respectively; **Fig. 8**). However, only reductions for the lactonase treatment at the two different sampling depths were statistically significant. These results reinforce the bacterial order analyses above, which demonstrated that SRB relative abundance was correlated with corrosion. The decrease of *dsrA* gene abundance in lactonase-treated samples suggests that a lactonase additive can reduce the abundance of SRBs on a steel surface. This observation is interesting and consistent with previous observations on the SRB *Desulfovibrio vulgaris*, for which QS inhibitors reduced biofilm and biocorrosion rates, while AHL signals increased these parameters [64]. In our work reported here, both direct (SRB metabolism) and indirect (reduction in abundance) effects might be at play to explain the observed reduction in biocorrosion.

**Fig 8.**
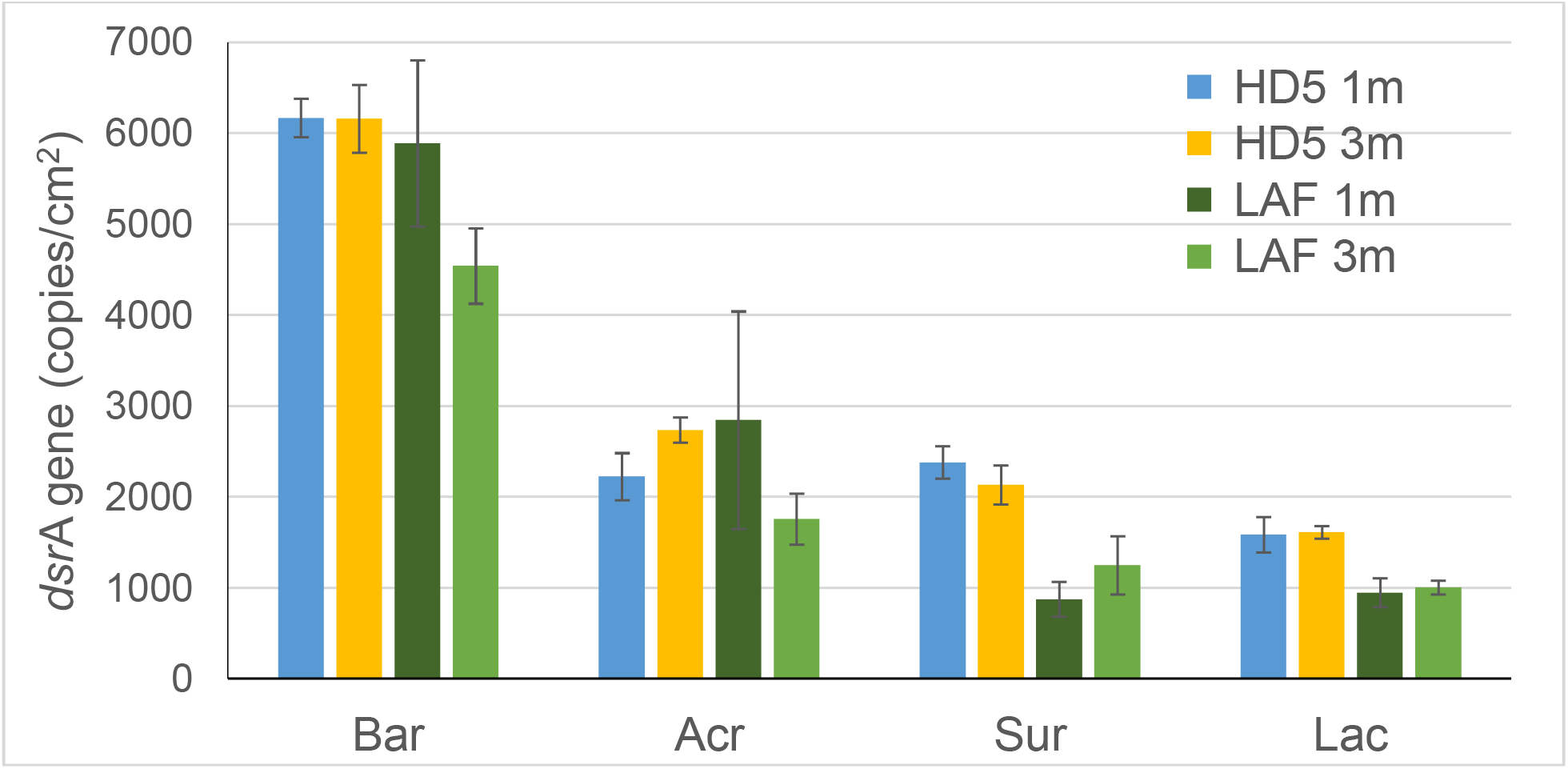
Estimation of SRBs using qPCR of an indicator gene (*dsr*A). *dsr*A gene (copies/cm^2^) in DNA extracted on steel coupons with different experimental treatments after 8 months of exposure. Bar: Bare steel control without coating, Acr: Acrylic coating control, Sur: Acrylic coating with 200 ug/ml surfactin, Lac: Acrylic coating with 200 ug/ml SsoPox lactonase enzyme. Mean values of duplicate coupons are shown (n=2).

Taken together, ours results suggest that corrosion reduction can be caused by subtle differences in bacterial community composition, including SRB and IOB, within surface biofilms and corrosion tubercles, even if they do not provide a definitive mechanism for the observed corrosion reduction by quorum quenching lactonase enzymes. The concurrent reduction in tubercle formation and change in the bacterial community composition on lactonase-treated coupons confirmed that steel corrosion can be altered in the field by modifying attached bacterial communities.

## Conclusion

The potential of a stable quorum quenching lactonase as an anticorrosion coating additive was evaluated in a field study. Over the course of 21 months of submersion in two sites and at two different depths, the non-biocidal enzymatic coating outperformed controls, including the biological biocide surfactin, in reducing corrosion on steel samples in the Duluth-Superior Harbor. This provides evidence for the potential of quorum quenching enzymes as viable antifouling coating additives. Remarkably, surface microbial population analysis revealed very minor changes in the microbial communities on all of the steel coupons. This finding indicated that biologically-influenced corrosion, a very significant phenomenon in the Duluth-Superior Harbor, may be due to groups of bacteria present at the surface in limited abundance. A detailed analysis indicated that key changes in surface bacterial communities correlated with corrosion levels, and may partly explain the observed differences in corrosion levels. In particular, sulfate-reducing bacteria (SRBs) appeared were less common or their genes (dsrA) reduced on the surface of lactonase-coated coupons. Future studies may provide more insights into the specific mechanisms of the reduction in bio-induced corrosion by quorum quenching enzymes. Such knowledge may allow specific targeting of key microbial groups while minimizing effects on other, unrelated environmental bacteria.

## Acknowledgement

Funding for this project was partly provided by the Minnesota Environment and Natural Resources Trust Fund as recommended by the Minnesota Aquatic Invasive Species Research Center (MAISRC) and the Legislative-Citizen Commission on Minnesota Resources (LCCMR). Other funding was provided by the University of Minnesota MnDRIVE initiative. This work was also sponsored by the Minnesota Sea Grant College Program to MHE and REH supported by the NOAA office of Sea Grant, United States Department of Commerce, under grant No. NA18OAR417010. The statements, findings, conclusions, and recommendations are those of the author(s) and do not necessarily reflect the views of NOAA, the Sea Grant College Program, or the U.S. Department of Commerce. The U.S. Government is authorized to reproduce and distribute reprints for government purposes, notwithstanding any copyright notation that may appear hereon. This paper is journal reprint No. JR 697 of the Minnesota Sea Grant College Program. The authors wish to thank AMI Consulting Engineering for providing, placing and retrieving steel coupons in the Duluth-Superior harbor.

