## Supplementary material for "Field testing of an enzymatic quorum quencher coating additive to reduce biocorrosion of steel"

### Summary

Fig S13. Nonmetric multidimensional scaling plot showing the differences between bacterial communities in different treatments on corroding steel coupons grouped by site and sample exposure time. ........................................ Erreur ! Signet non défini.

**Table S1**. Analysis of molecular variance (AMOVA) showing the genetic variation among different treatment groups, sampling times, sites, and depths.

| Compared parameter | Compared samples | AMOVA FST (%) | p-value |
|---|---|---|---|
| TREATMENTS | **Acrylic_Control-** | 2.92137 | 0.003* |
| | **Bare_Steel_Control-** | | |
| | **all samples** | | |
| | **Acrylic_Control-** | 1.50319 | 0.213 |
| | **Lactonase-all samples** | 3.45146 | 0.164 |
| | 1 month-HD5-1m | 1.23856 | 0.436 |
| | 1 month-HD5-3m | 1.86708 | 0.134 |
| | 1 month-LAF-1m | 0.78288 | 0.335 |
| | 1 month-LAF-3m | 1.33684 | 0.147 |
| | 2 months-HD5-1m | 0.71462 | 0.45 |
| | 2 months-HD5-3m | 0.31260 | 0.733 |
| | 2 months-LAF-1m | 1.55857 | 0.214 |
| | 2 months-LAF-3m | 2.14143 | 0.281 |
| | 8 months-HD5-1m | 2.20429 | 0.368 |
| | 8 months-HD5-3m | 2.07265 | 0.312 |
| | 8 months-LAF-1m | 3.37534 | 0.202 |
| | 8 months-LAF-3m | 4.66278 | 0.105 |
| | 21 months-HD5-1m | 3.29742 | 0.066 |
| | 21 months-HD5-3m | 3.62930 | 0.128 |
| | 21 months-LAF-1m | 4.24146 | 0.074 |
| | 21 months-LAF-3m | | |
| | **Acrylic_Control-** | 1.02473 | 0.409 |
| | **Surfactin-all samples** | 3.10409 | 0.188 |
| | 1 month-HD5-1m | 1.18736 | 0.496 |
| | 1 month-HD5-3m | 1.92413 | 0.142 |
| | 1 month-LAF-1m | 2.14667 | 0.125 |
| | 1 month-LAF-3m | 0.41522 | 0.526 |
| | 2 months-HD5-1m | 0.62950 | 0.689 |
| | 2 months-HD5-3m | 0.25412 | 0.740 |
| | 2 months-LAF-1m | 2.05739 | 0.196 |
| | 2 months-LAF-3m | 2.57217 | 0.263 |
| | 8 months-HD5-1m | 2.97855 | 0.185 |
| | 8 months-HD5-3m | 3.47653 | 0.156 |
| | 8 months-LAF-1m | 3.36844 | 0.180 |
| | 8 months-LAF-3m | 2.96327 | 0.206 |
| | 21 months-HD5-1m | 1.83590 | 0.178 |
| | 21 months-HD5-3m | 5.96721 | 0.055 |
| | 21 months-LAF-1m | 3.85772 | 0.096 |
| | 21 months-LAF-3m | | |
| | **Bare_Steel_Control-** | 5.87454 | <0.001* |
| | **Lactonase-all samples** | | |
| | **Bare_Steel_Control-** | 4.23742 | <0.001* |
| | **Surfactin-all samples** | | |
| | **Lactonase-Surfactin-** | 0.95457 | 0.467 |
| | **all samples** | 0.13241 | 0.872 |
| | 1 month-HD5-1m | 0.21687 | 0.769 |

| | | | |
|---|---|---|---|
| | 1 month-HD5-3m<br>1 month-LAF-1m<br>1 month-LAF-3m<br>2 months-HD5-1m<br>2 months-HD5-3m<br>2 months-LAF-1m<br>2 months-LAF-3m<br>8 months-HD5-1m<br>8 months-HD5-3m<br>8 months-LAF-1m<br>8 months-LAF-3m<br>21 months-HD5-1m<br>21 months-HD5-3m<br>21 months-LAF-1m<br>21 months-LAF-3m | 0.24552<br>0.89360<br>1.21166<br>0.46538<br>0.28542<br>1.26277<br>1.89201<br>2.54260<br>2.18812<br>1.02574<br>3.17190<br>2.91945<br>6.41289<br>0.96621 | 0.624<br>0.335<br>0.217<br>0.640<br>0.795<br>0.266<br>0.311<br>0.283<br>0.188<br>0.213<br>0.107<br>0.084<br>0.053<br>0.642 |
| EXPOSURE PERIOD | 1 Month – 2 Months | 59.2133 | <0.001* |
| | 1 Month - 21 Months | 32.8885 | <0.001* |
| | 1 Month - 8 Months | 35.0827 | <0.001* |
| | 2 Months -21 Months | 47.9193 | <0.001* |
| | 2 Months -8 Months | 54.6192 | <0.001* |
| | 21 Months -8 Months | 3.22321 | 0.007* |
| SITE | HD5-LAF | 2.42668 | 0.02* |
| DEPTH | 1 M-3 M | 5.5809 | <0.001* |

**Table S2**. Summary of AHL-producing, AHL-degrading, and AHL-sensing ability of the bacterial orders of highest relative abundance.

| ORDERS OF HIGHEST ABUNDANCE | TYPE | AHL PRODUCING | AHL DEGRADING | AHL SENSING | REFERENCES |
|---|---|---|---|---|---|
| *BDELLOVIBRIONALES* | SRB | | | * | [1] |
| *DESULFOVIBRIONALES* | SRB | * | * | * | [2-6] |
| *DESULFUROMONADALES* | SRB | * | | * | [2, 7-8] |
| *METHYLOCOCCALES* | SRB | * | | * | [9-10] |
| *METHYLOPHILALES* | SRB | * | | * | [10] |
| *SULFURICELLALES* | SRB | | | * | [11] |
| *VERRUCOMICROBIALES* | SRB | | | * | [10,12] |
| *ACIDIMICROBIALES* | IOB | * | * | * | [10] |
| *ACTINOMYCETALES* | IOB | * | * | * | [13-14] |
| *BACTEROIDALES* | IOB | * | | * | [15] |
| *BURKHOLDERIALES* | IOB | * | | * | [10, 16-17] |
| *FLAVOBACTERIALES* | IOB | | * | * | [10, 18] |
| *GALLIONELLALES* | IOB | | | * | [19] |
| *HOLOPHAGALES* | IOB | | | * | [12] |
| *NITROSOMONADALES* | IOB | * | | * | [20-22] |
| *NITROSPIRALES* | IOB | * | * | * | [10, 23] |
| *PLANCTOMYCETALES* | IOB | * | * | * | [24] |
| *RHIZOBIALES* | IOB | * | * | * | [10,18,25] |
| *RHODOBACTERALES* | IOB | * | * | * | [10,18,25-26] |
| *RHODOCYCLALES* | IOB | * | | * | [27-28] |
| *RHODOSPIRILLALES* | IOB | * | * | * | [10,29-30] |
| *SPHINGOBACTERIALES* | IOB | * | * | * | [10,18,31-32] |
| *SPHINGOMONADALES* | IOB | * | * | * | [33-34] |
| *ACIDOBACTERIA* | others | * | | * | [23,35] |
| *ARIDIBACTER* | others | | | | ⸸ |
| *BACILLALES* | others | * | * | * | [36-37] |
| *CAULOBACTERALES* | others | | | * | [38] |
| *CYTOPHAGALES* | others | | * | | [10, 39] |
| *GAIELLALES* | others | | | | ⸸ |
| *GEMMATIMONADALES* | others | | * | | [10] |
| *MYXOCOCCALES* | others | * | * | * | [10, 40-41] |
| *OPITUTALES* | others | | * | | [10] |
| *PSEUDOMONADALES* | others | * | * | * | [42-44] |
| *RICKETTSIALES* | others | * | | * | [45-46] |
| *SACCHARIBACTERIA* | others | * | | * | [47-49] |

| | | | | | |
|---|---|---|---|---|---|
| *XANTHOMONADALES* | others | | * | * | [27, 50-51] |

ǂ information could not be found in the literature.

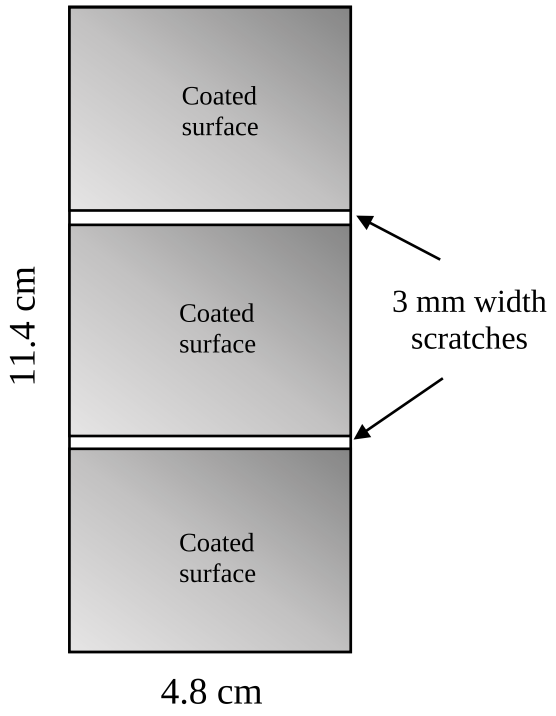

**Fig S1**. Schematic of a coating treated steel coupon and the scratches.

| | A: No coating control | B: Acrylic coating control | | C: 200 ug/ml surfactin | | D: 200 ug/ml lactonase | |
|---|---|---|---|---|---|---|---|
| | | Unscratched | Scratched | Unscratched | Scratched | Unscratched | Scratched |
| 1 month | 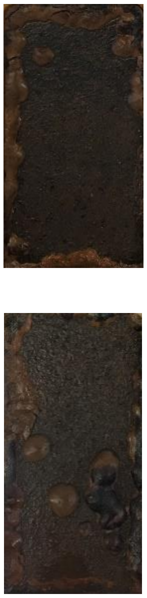 | 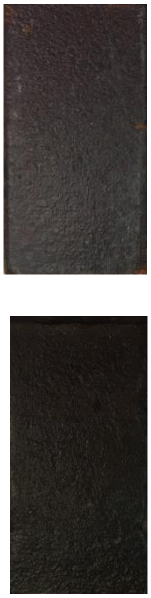 | 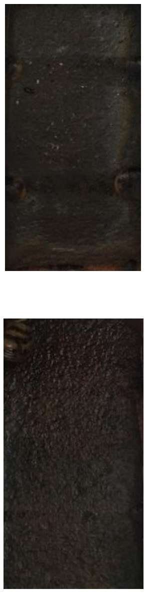 | 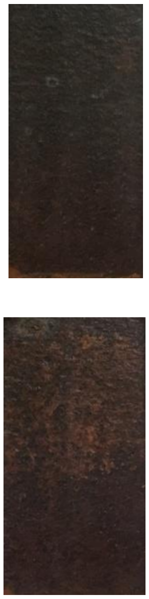 | 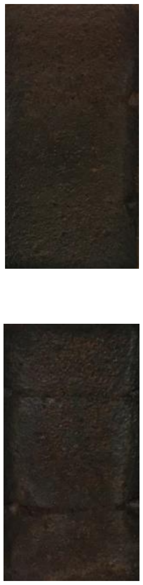 | 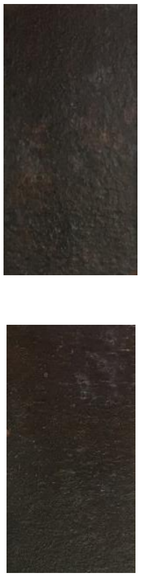 | 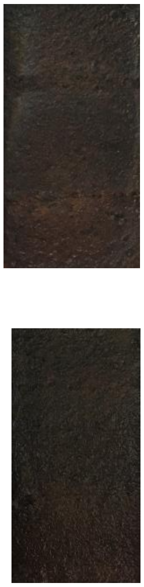 |
| 2 months | 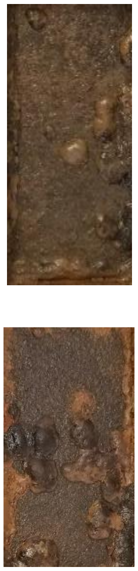 | 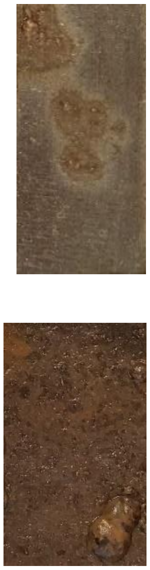 | 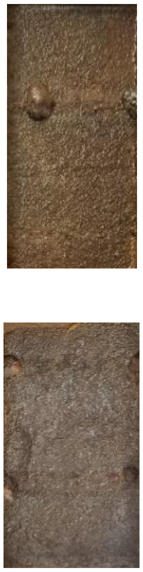 | 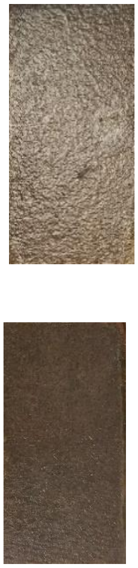 | 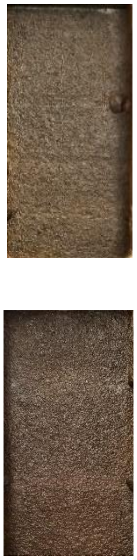 | 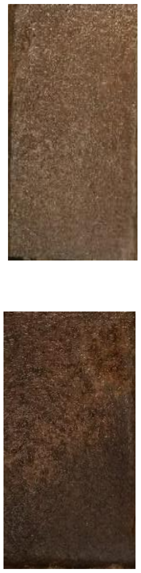 | 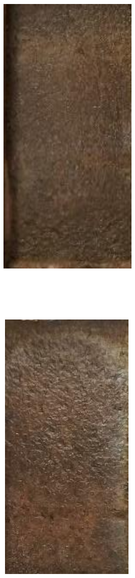 |

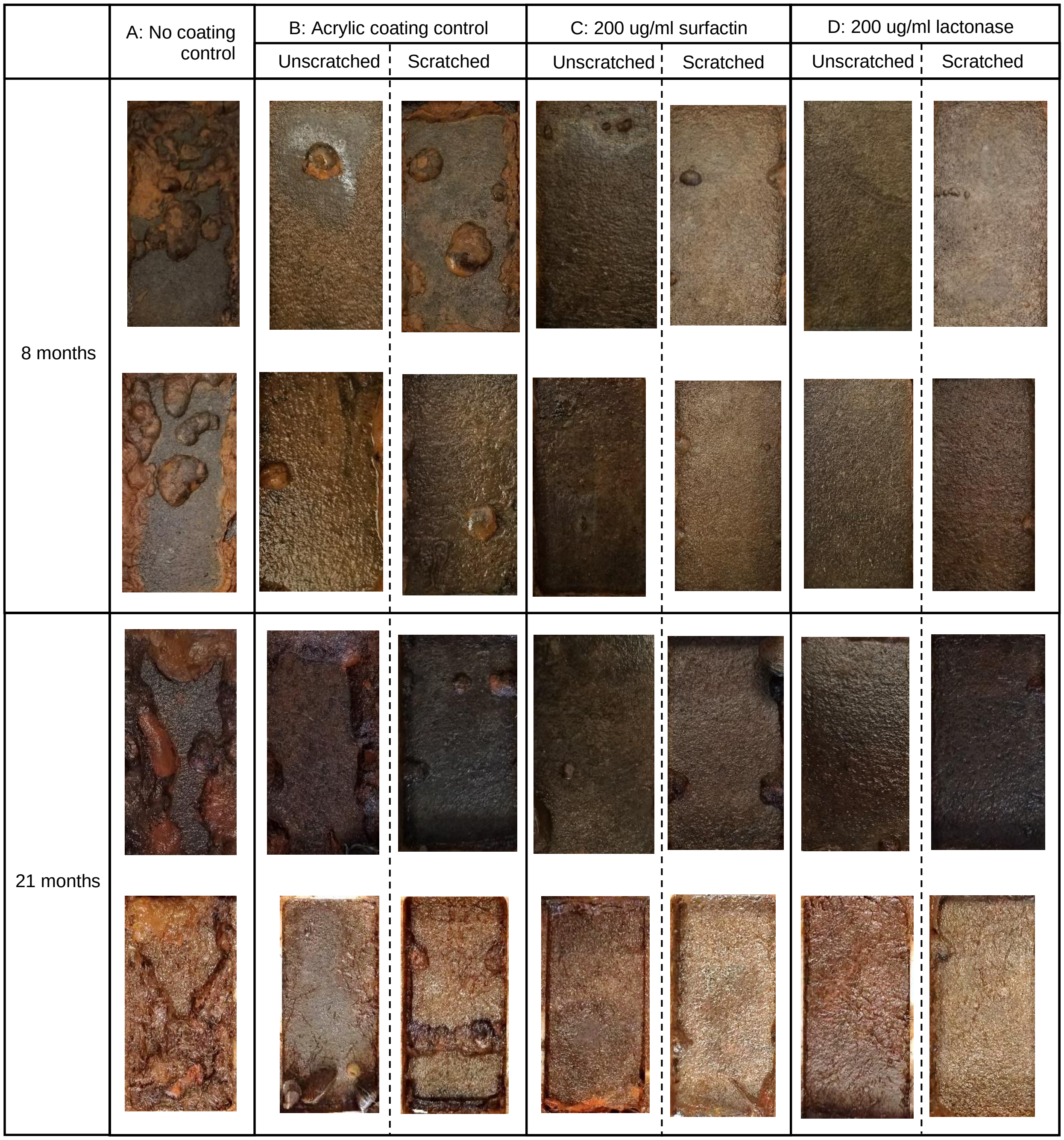

**Fig S2**. Photographs of control and treated steel coupons after 1, 2, 8 and 21 months of exposure in the Duluth-Superior Harbor (Site: HD5, Depth: 3m). This set of pictures is representative of sample coupons at other sites/depth. The image on the left in each test group is the unscratched side of the coupon.

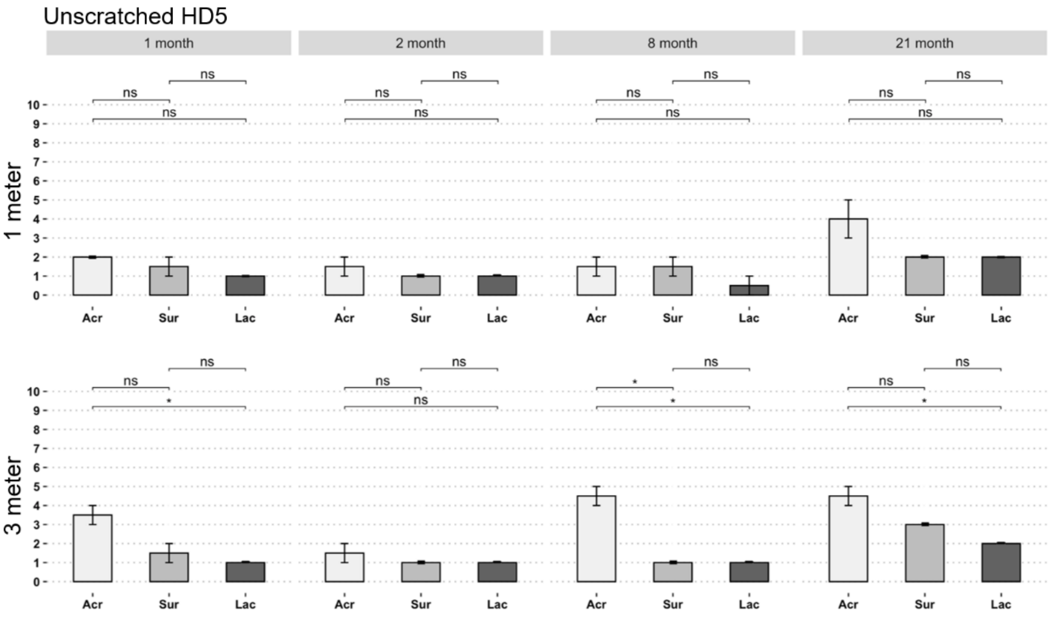
Unscratched HD5
1 month
2 month
8 month
21 month
1 meter
3 meter
Acr
Sur
Lac
ns
*

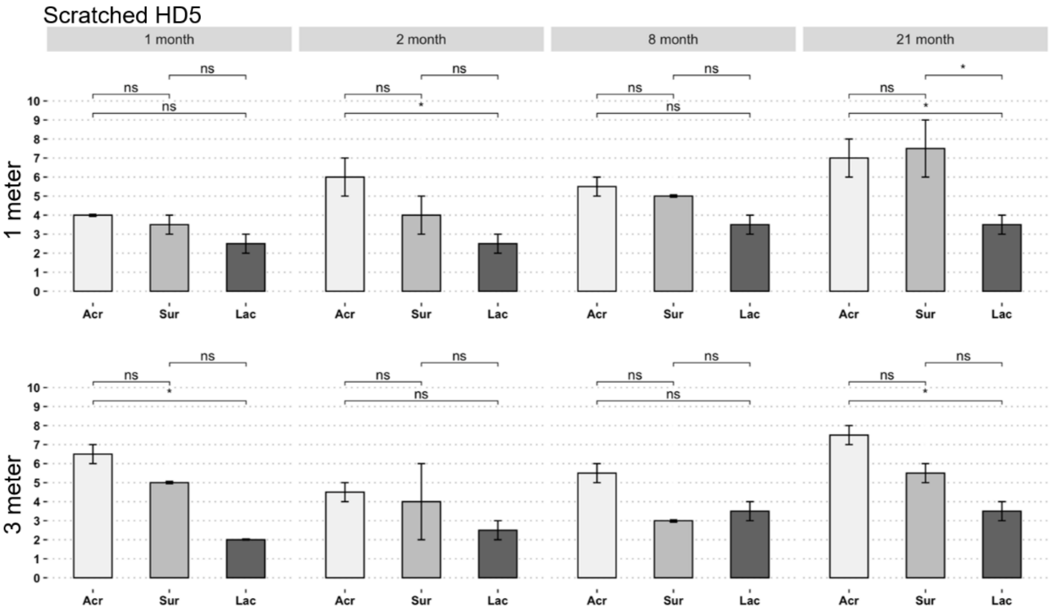
Scratched HD5
1 month
2 month
8 month
21 month
1 meter
3 meter
Acr
Sur
Lac
ns
*

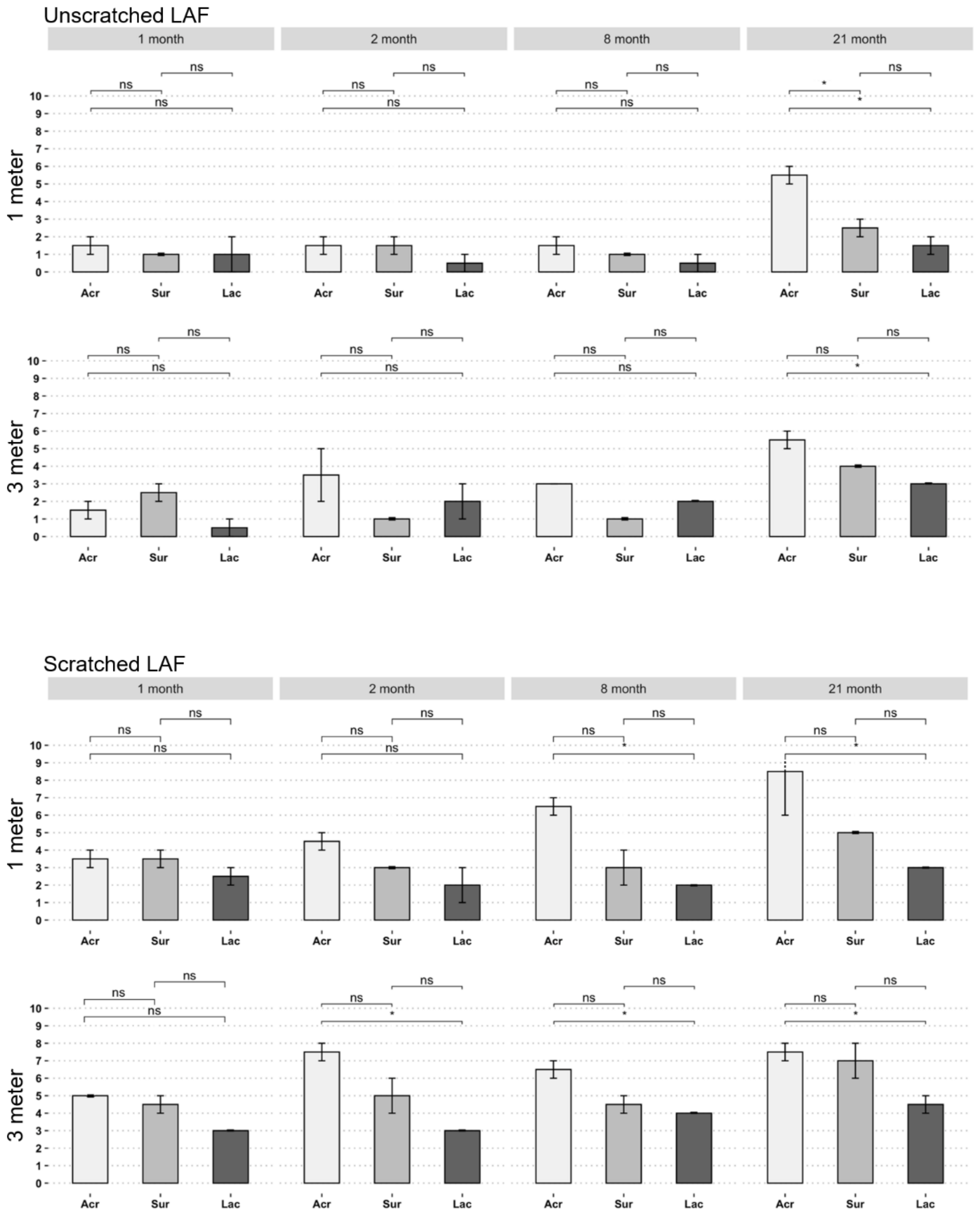

**Fig S3**. Counts of corrosion tubercles on the duplicate unscratched and scratched steel coupons with different experimental treatments at the two tested sites (HD5 and LAF) and depths (1 and 3m). Legend - Acr: Acrylic coating control, Sur: Acrylic coating with 200 ug/ml surfactin, Lac: Acrylic coating with 200 ug/ml *Sso*Pox lactonase enzyme.

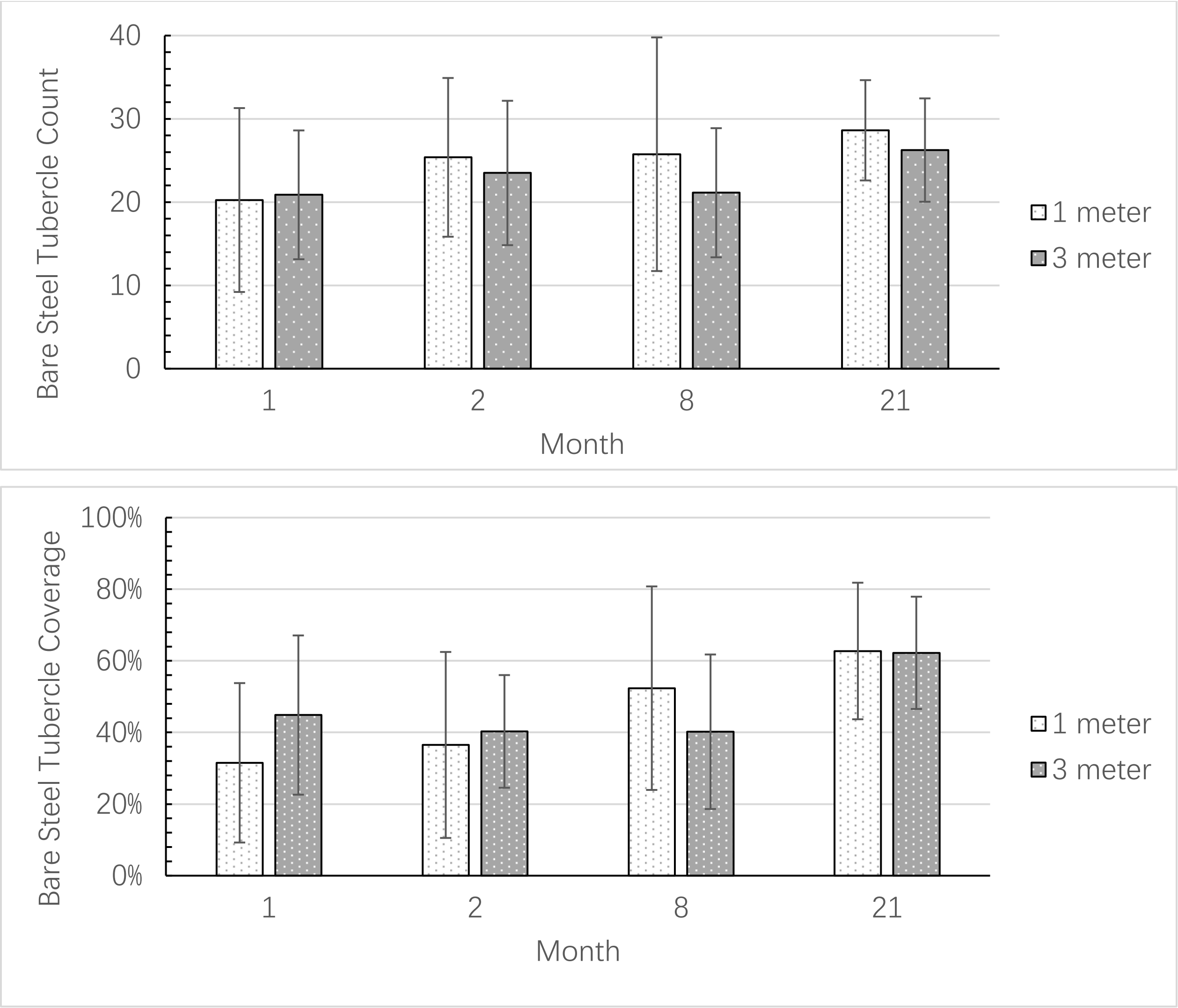

**Fig S4**. Tubercle counts (upper panel) and coverage (lower panel) of corrosion tubercles on bare steel coupons at 4 sampling times. Mean values for 2 coupons at both sites are shown (n=4).

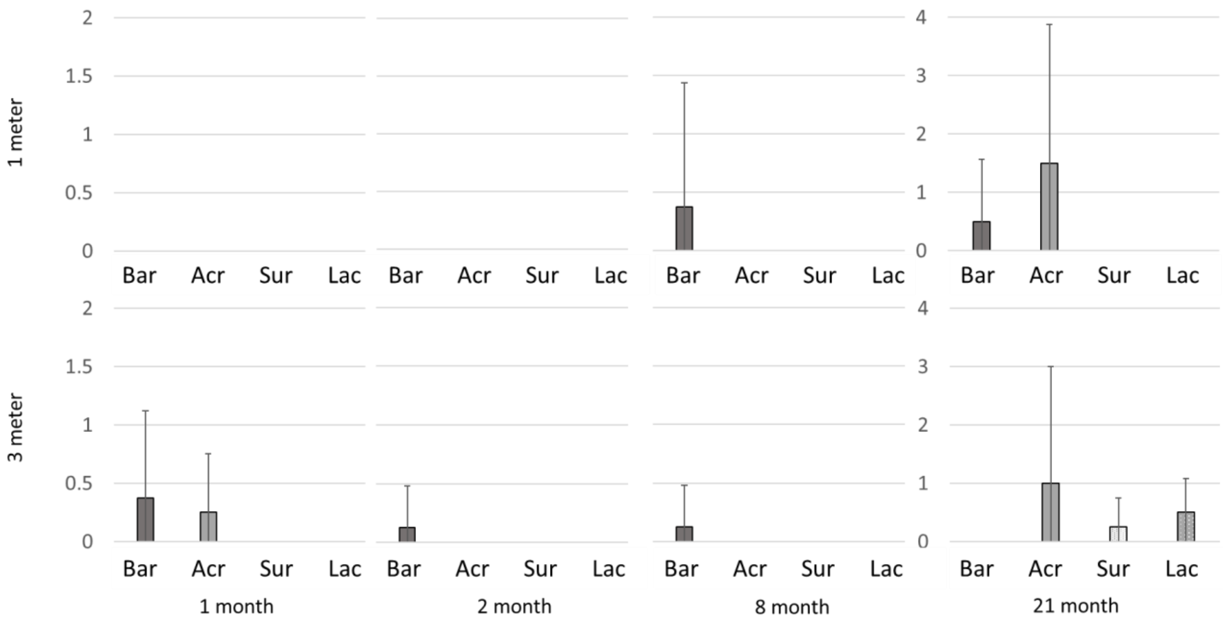

**Fig S5**. Mussel counts on unscratched steel coupons from different experimental treatments. Mean values for 2 coupons at both sites are shown (n=4). Legend - Bar: Bare Steel, Acr: Acrylic coating control, Sur: Acrylic coating with 200 ug/ml surfactin, Lac: Acrylic coating with 200 ug/ml *Sso*Pox lactonase enzyme.

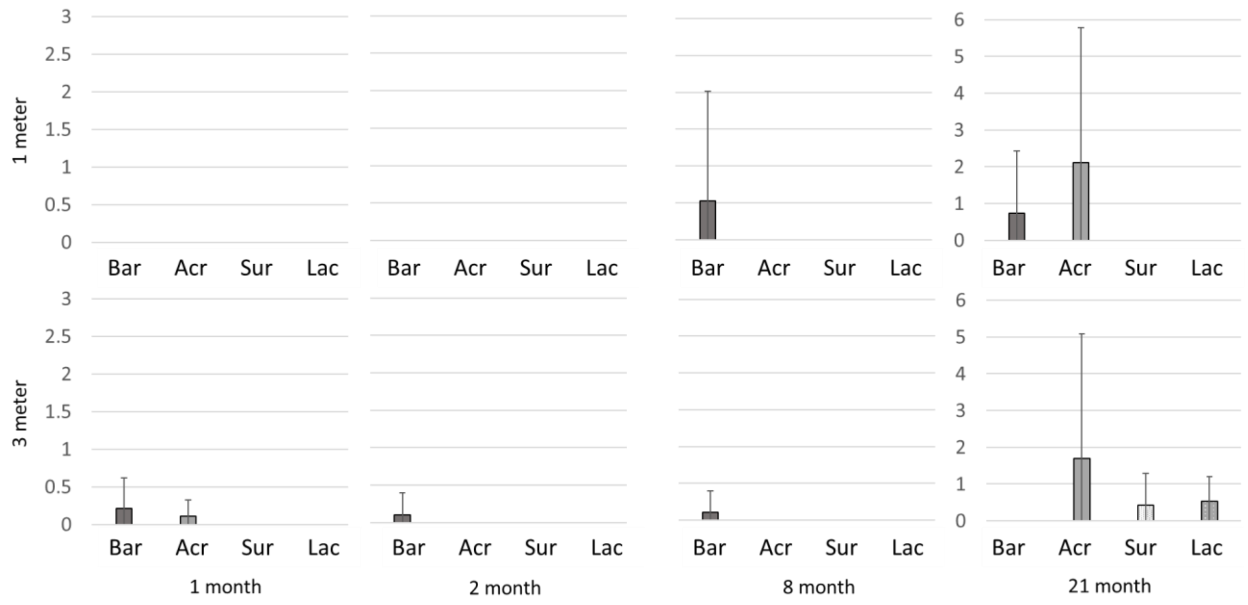

**Fig. S6**. Mussel weights on unscratched steel coupons from different experimental treatments. Mean values for 2 coupons at both sites are shown (n=4). Legend - Bar: Bare Steel, Acr: Acrylic coating control, Sur: Acrylic coating with 200 ug/ml surfactin, Lac: Acrylic coating with 200 ug/ml *Sso*Pox lactonase enzyme.

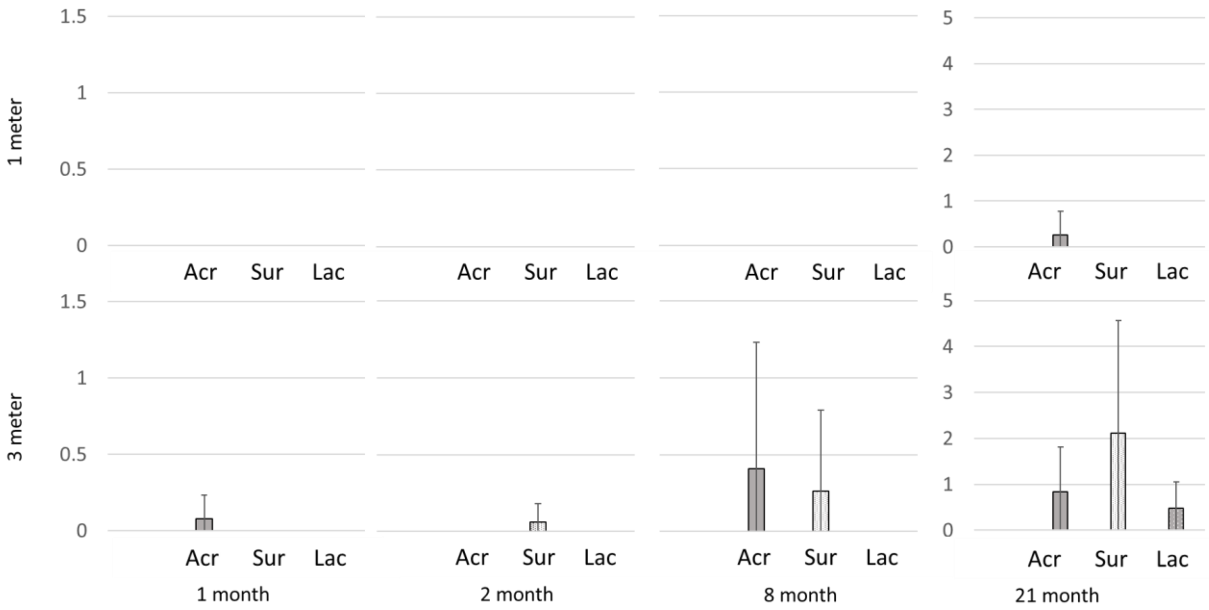

**Fig S7**. Mussel counts on scratched steel coupons from different experimental treatments. Mean values for 2 coupons at both sites are shown (n=4). Legend - Acr: Acrylic coating control, Sur: Acrylic coating with 200 ug/ml surfactin, Lac: Acrylic coating with 200 ug/ml *Sso*Pox lactonase enzyme.

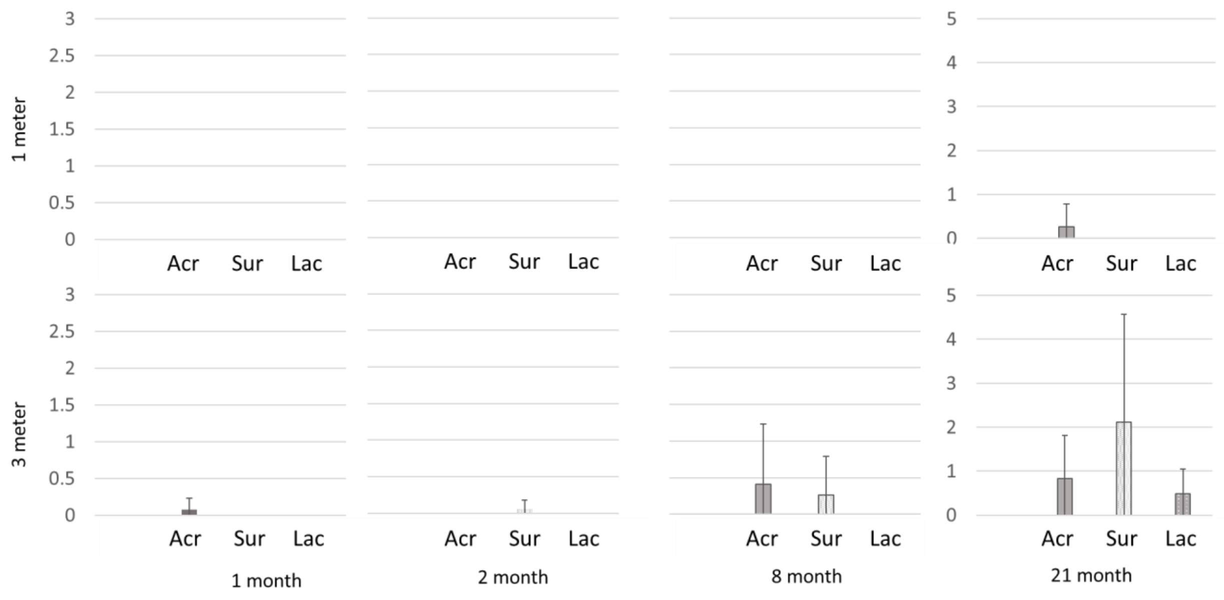

**Fig S8**. Mussel weights on scratched steel coupons from different experimental treatments. Mean values for 2 coupons at both sites are shown (n=4). Legend - Acr: Acrylic coating control, Sur: Acrylic coating with 200 ug/ml surfactin, Lac: Acrylic coating with 200 ug/ml *Sso*Pox lactonase enzyme.

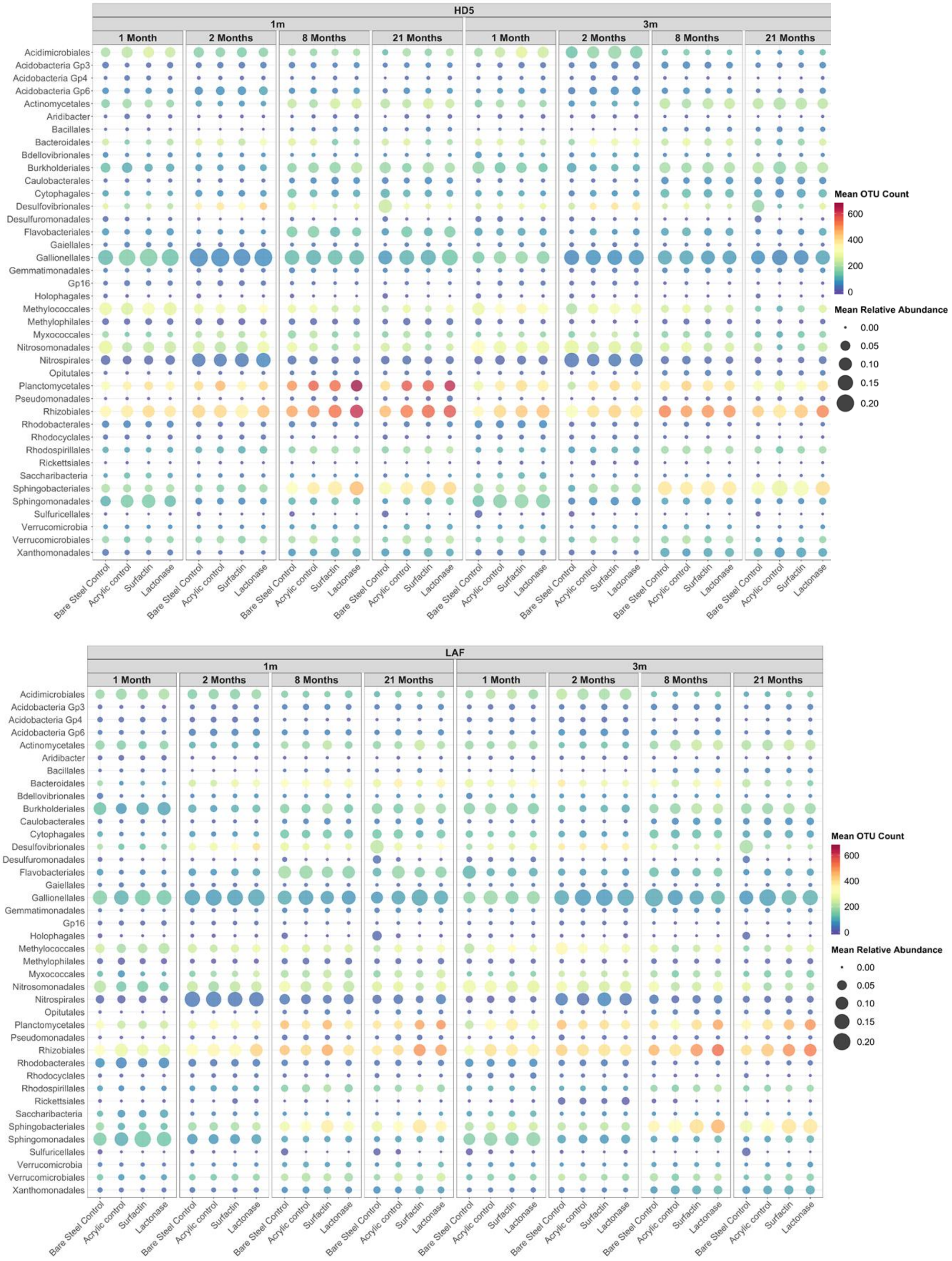

**Fig S9**. Heatmap comparing the relative abundance of partial 16S rRNA sequences and OTU richness for the top 40 bacterial orders in all samples from each treatment and control. Diversity is indicated by the number of OTUs in each bacterial order.

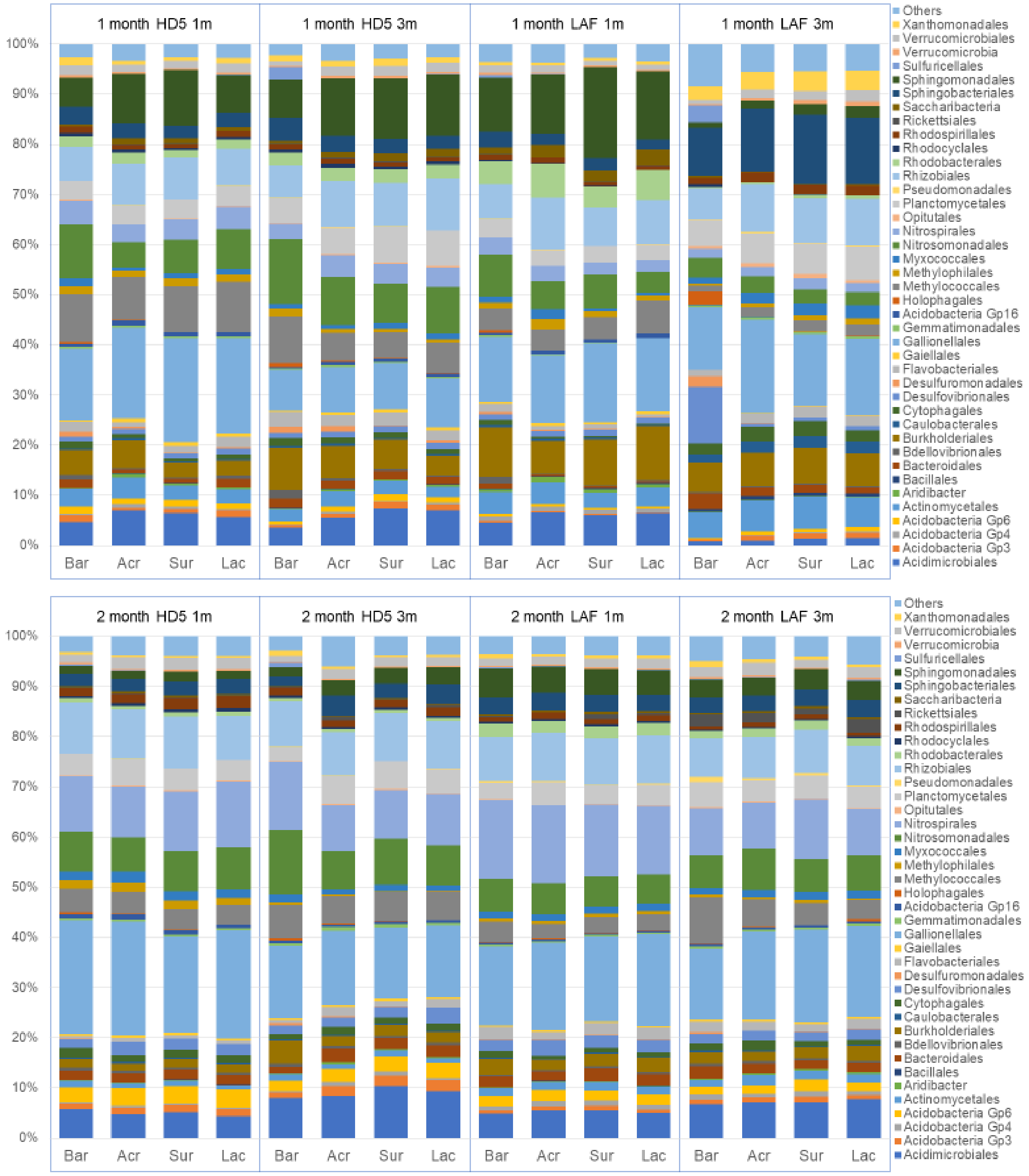
1 month HD5 1m
1 month HD5 3m
1 month LAF 1m
1 month LAF 3m
2 month HD5 1m
2 month HD5 3m
2 month LAF 1m
2 month LAF 3m
100%
90%
80%
70%
60%
50%
40%
30%
20%
10%
0%
Bar
Acr
Sur
Lac
Others
Xanthomonadales
Verrucomicrobiales
Verrucomicrobia
Sulfuricellales
Sphingomonadales
Sphingobacteriales
Saccharibacteria
Rickettsiales
Rhodospirillales
Rhodocyclales
Rhodobacterales
Rhizobiales
Pseudomonadales
Planctomycetales
Opitutales
Nitrospirales
Nitrosomonadales
Myxococcales
Methylophilales
Methylococcales
Holophagales
Acidobacteria Gp16
Gemmatimonadales
Gallionellales
Gaiellales
Flavobacteriales
Desulfuromonadales
Desulfovibrionales
Cytophagales
Caulobacterales
Burkholderiales
Bdellovibrionales
Bacteroidales
Bacillales
Aridibacter
Actinomycetales
Acidobacteria Gp6
Acidobacteria Gp4
Acidobacteria Gp3
Acidimicrobiales

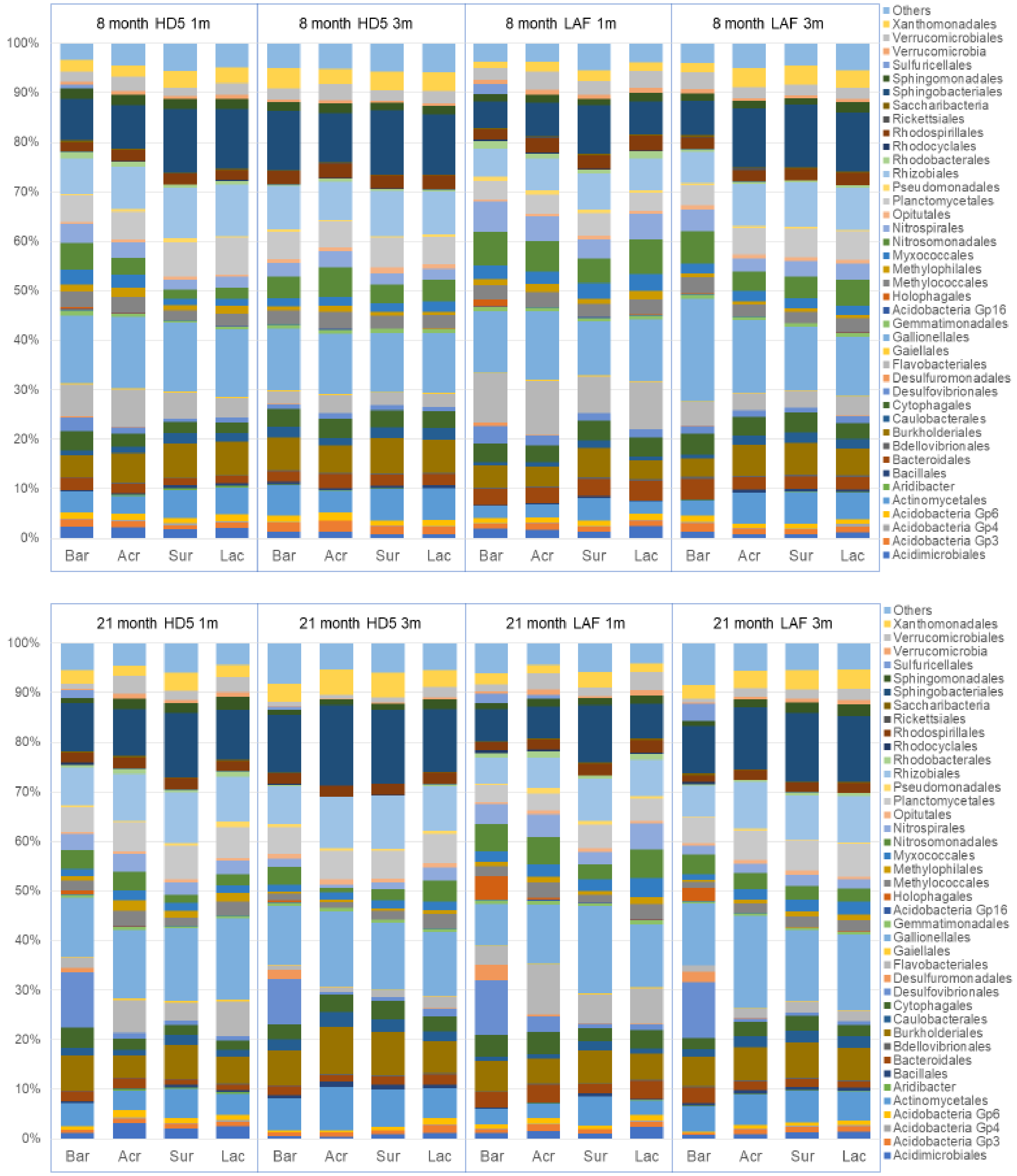

**Fig S10**. Relative abundance of partial 16S rRNA sequences for the top 40 bacterial orders of all samples from each treatment and control. Bar: Bare steel control without coating, Acr: Acrylic coating control, Sur: Acrylic coating with 200 ug/ml surfactin, Lac: Acrylic coating with 200 ug/ml SsoPox lactonase enzyme.

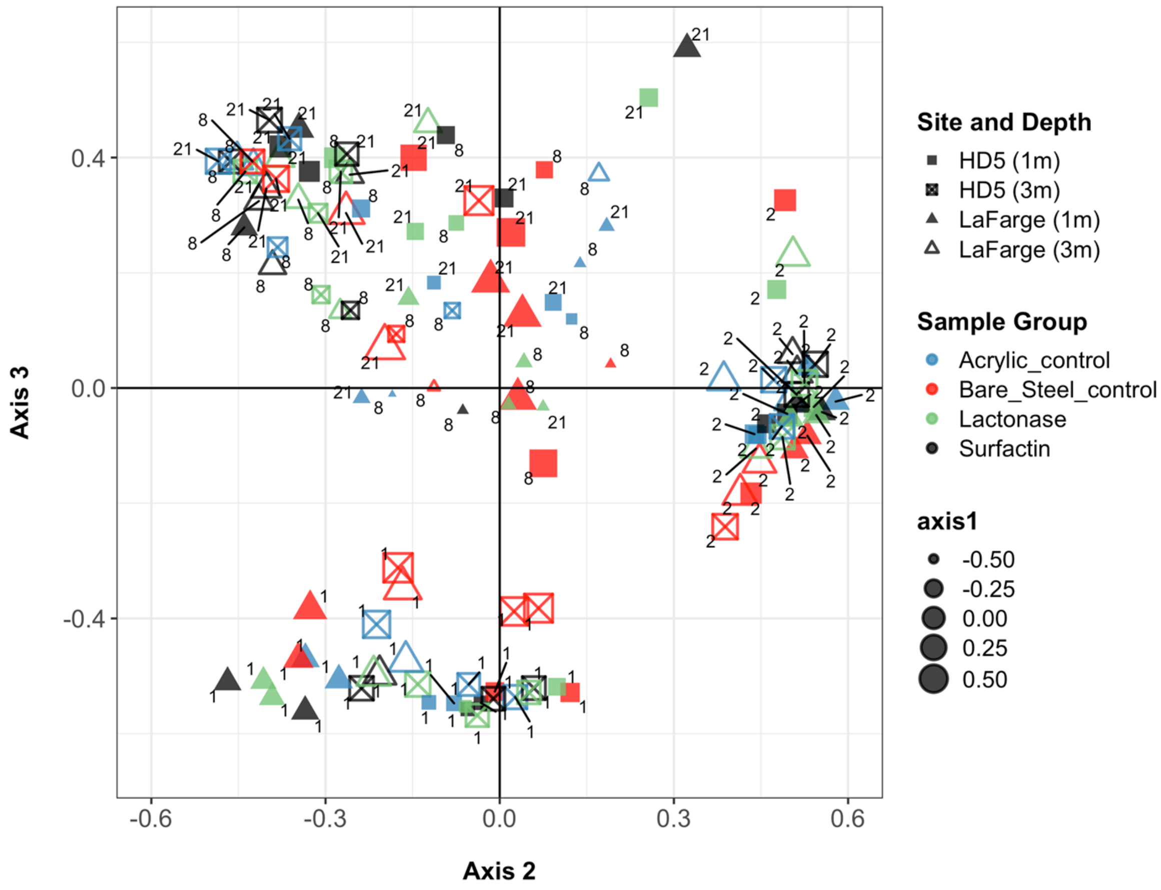

**Fig S11**. Nonmetric multidimensional scaling plot showing the differences between bacterial communities in different treatments on corroding steel coupons grouped by site and sample exposure time. The sample time of each sample is shown on the graph by number markings. The 4 treatment groups are separated by different colors, and the sample depths are separated by filled and hollow shapes. The stress value of this NMDS plot is lower than 0.1.

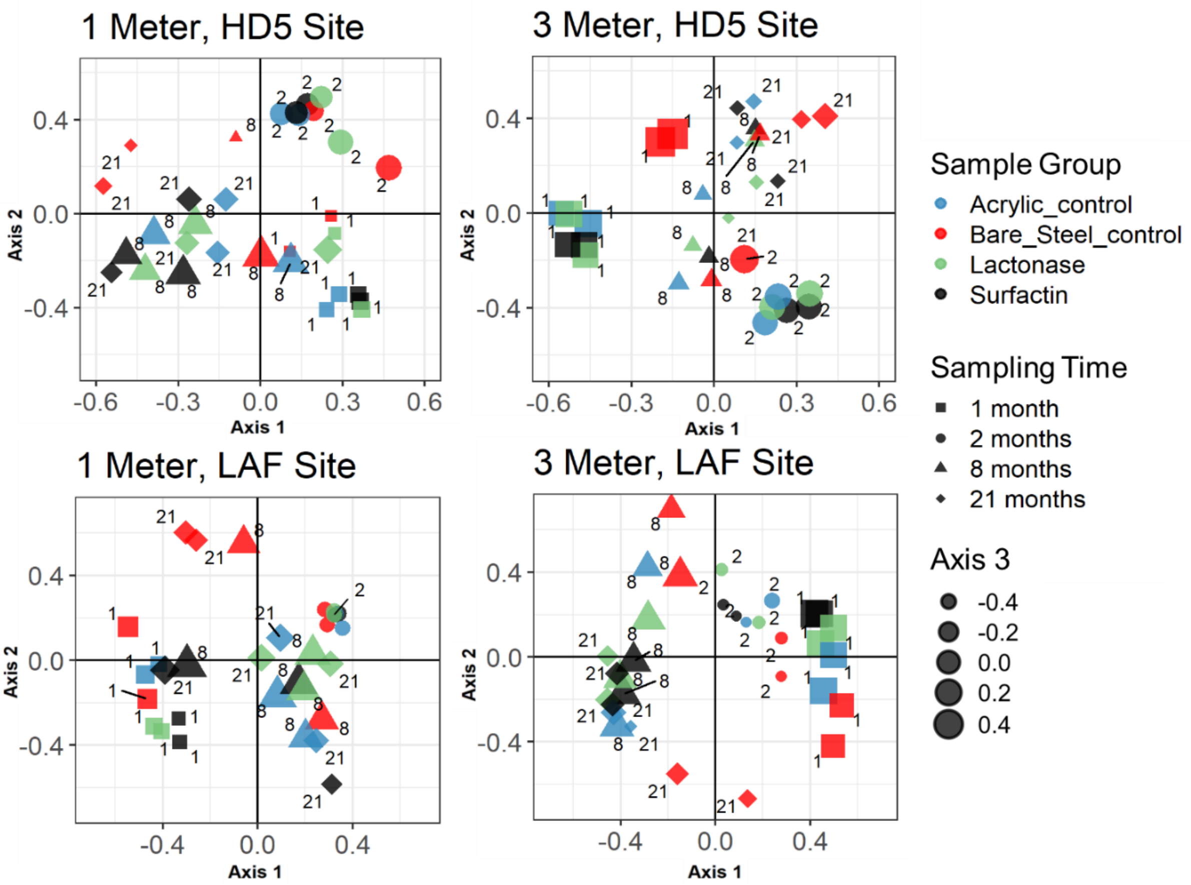

**Fig S12**. Nonmetric multidimensional scaling plot showing the differences between bacterial communities in different treatments on corroding steel coupons over time. The site and depth of each plot is shown above the graph. The 4 treatment groups are separated by different colors, and the sample months are separated by shapes. The stress value of each NMDS plot is lower than 0.1. NMDS plots of all data combined and of each site, depth and month are shown in Fig S10.

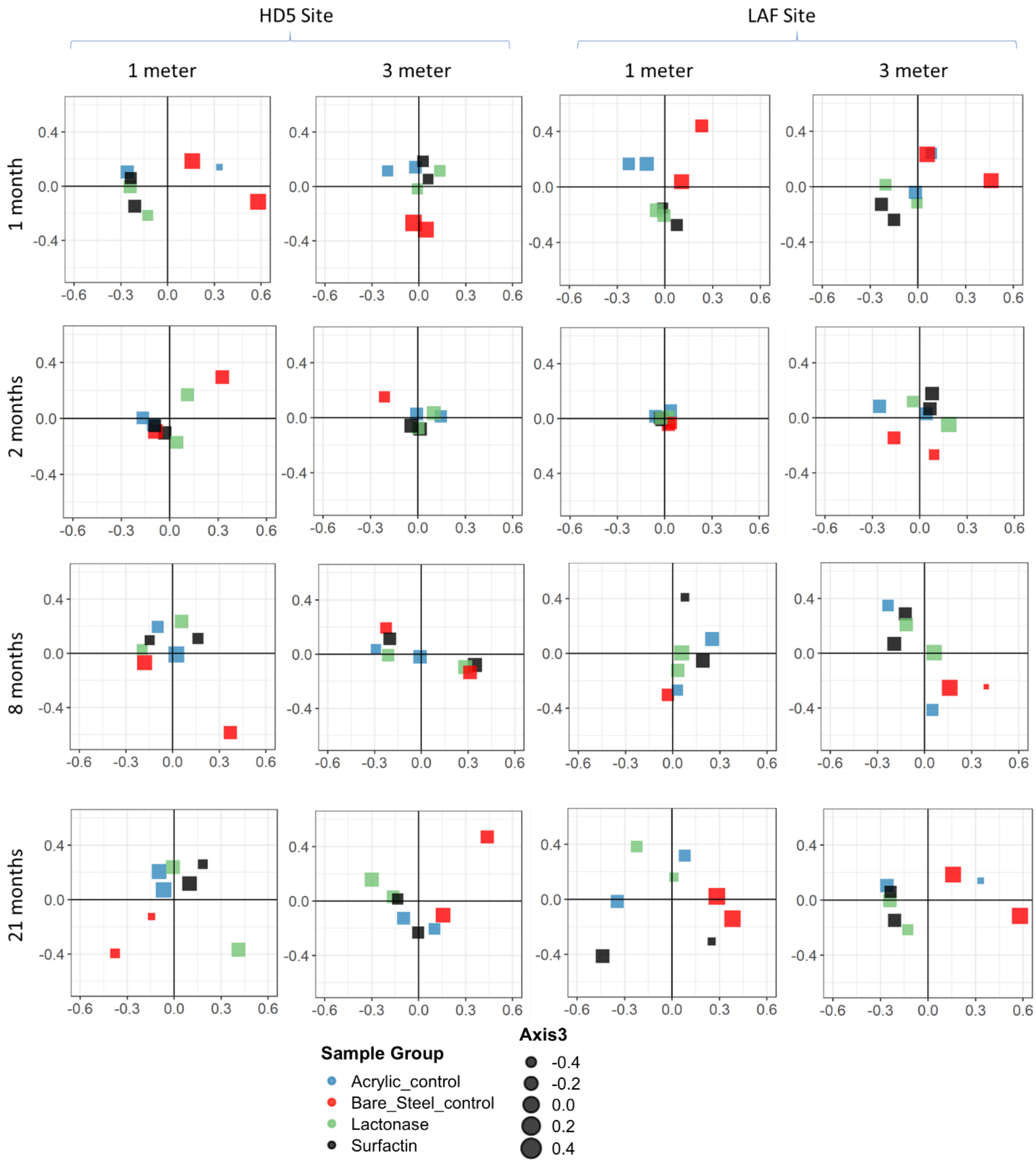

**Fig S13**. Nonmetric multidimensional scaling plot showing the differences between bacterial communities in different treatments on corroding steel coupons grouped by site and sample exposure time. The site, depth and months of each plot is shown above the graph. The 4 treatment groups are separated by different colors. The stress value of each NMDS plot is lower than 0.2.

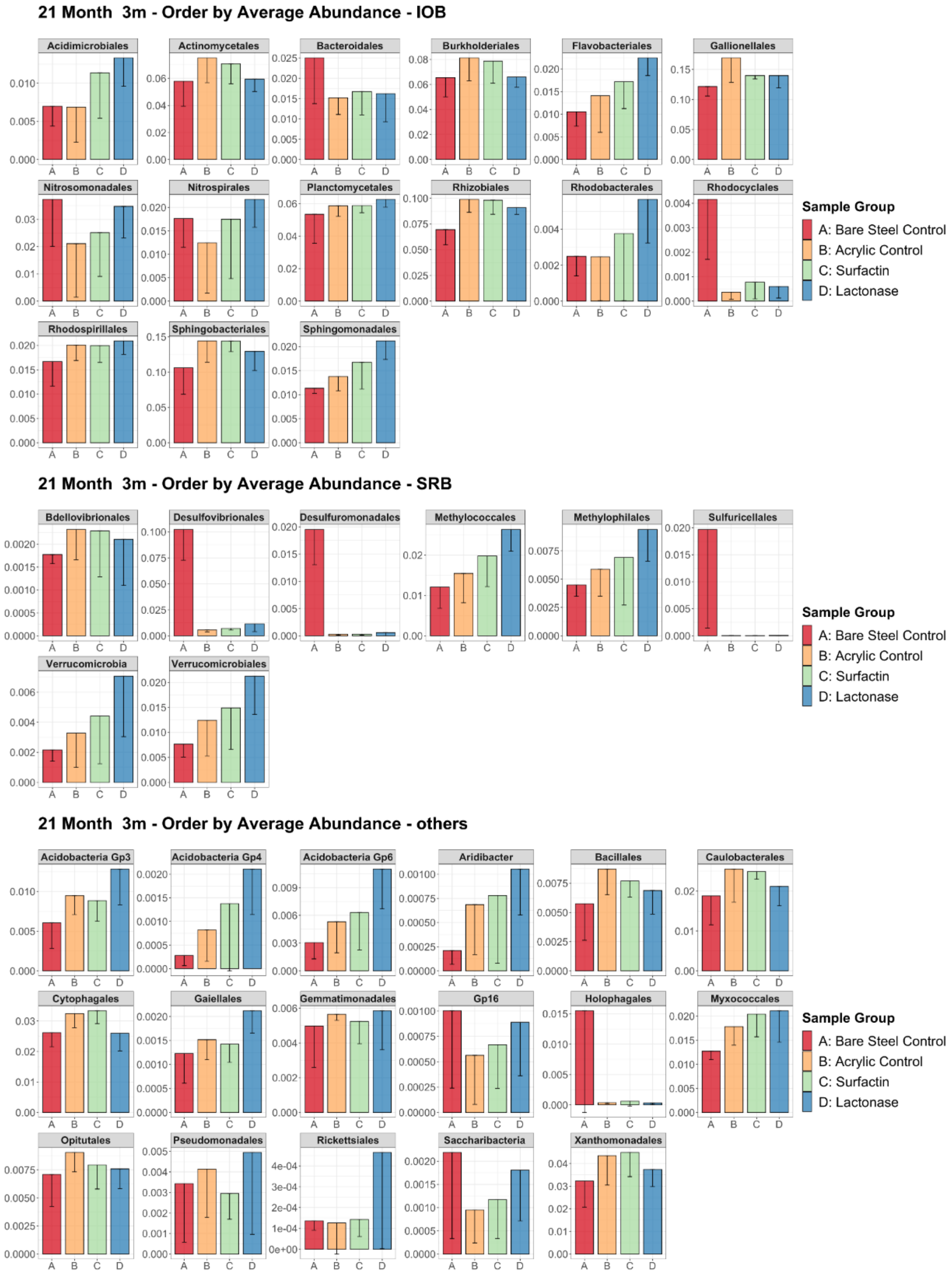

**Fig S14**. Average iron-oxidizing and sulfate-reducing bacteria relative abundances on steel coupons with different experimental treatments. A: Bare steel control without coating, B: Acrylic coating control, C: Acrylic coating with 200 ug/ml surfactin, D: Acrylic coating with 200 ug/ml *Sso*Pox lactonase enzyme. Mean values of 2 coupons of 21-month 3m at 2 sites are shown (n=4).

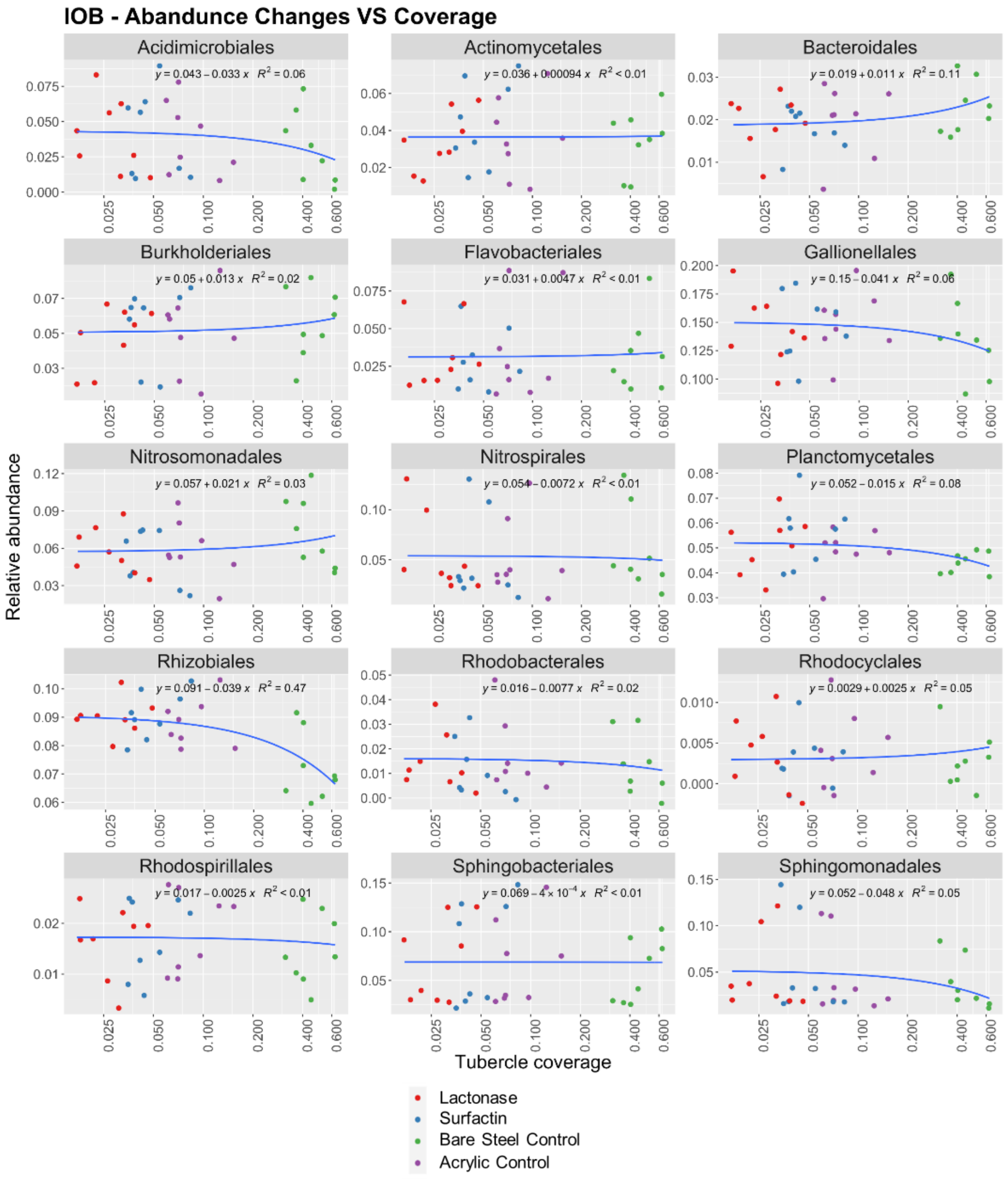
IOB - Abandunce Changes VS Coverage
Acidimicrobiales
y = 0.043 − 0.033 x  R² = 0.06
Actinomycetales
y = 0.036 + 0.00094 x  R² < 0.01
Bacteroidales
y = 0.019 + 0.011 x  R² = 0.11
Burkholderiales
y = 0.05 + 0.013 x  R² = 0.02
Flavobacteriales
y = 0.031 + 0.0047 x  R² < 0.01
Gallionellales
y = 0.15 − 0.041 x  R² = 0.06
Nitrosomonadales
y = 0.057 + 0.021 x  R² = 0.03
Nitrospirales
y = 0.054 − 0.0072 x  R² < 0.01
Planctomycetales
y = 0.052 − 0.015 x  R² = 0.08
Rhizobiales
y = 0.091 − 0.039 x  R² = 0.47
Rhodobacterales
y = 0.016 − 0.0077 x  R² = 0.02
Rhodocyclales
y = 0.0029 + 0.0025 x  R² = 0.05
Rhodospirillales
y = 0.017 − 0.0025 x  R² < 0.01
Sphingobacteriales
y = 0.069 − 4 × 10⁻⁴ x  R² < 0.01
Sphingomonadales
y = 0.052 − 0.048 x  R² = 0.05
Relative abundance
Tubercle coverage
Lactonase
Surfactin
Bare Steel Control
Acrylic Control

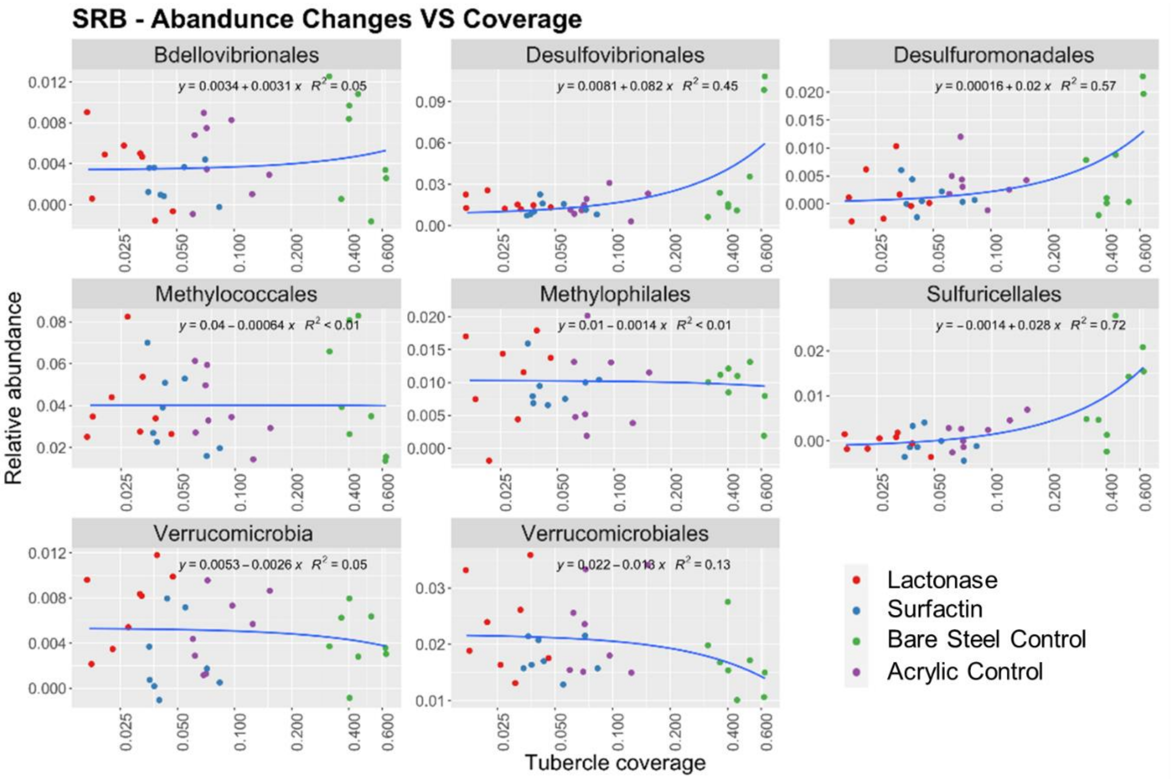
SRB - Abandunce Changes VS Coverage
Bdellovibrionales
y = 0.0034 + 0.0031 x  R^2 = 0.05
Desulfovibrionales
y = 0.0081 + 0.082 x  R^2 = 0.45
Desulfuromonadales
y = 0.00016 + 0.02 x  R^2 = 0.57
Methylococcales
y = 0.04 − 0.00064 x  R^2 < 0.01
Methylophilales
y = 0.01 − 0.0014 x  R^2 < 0.01
Sulfuricellales
y = −0.0014 + 0.028 x  R^2 = 0.72
Verrucomicrobia
y = 0.0053 − 0.0026 x  R^2 = 0.05
Verrucomicrobiales
y = 0.022 − 0.018 x  R^2 = 0.13
Relative abundance
Tubercle coverage
Lactonase
Surfactin
Bare Steel Control
Acrylic Control

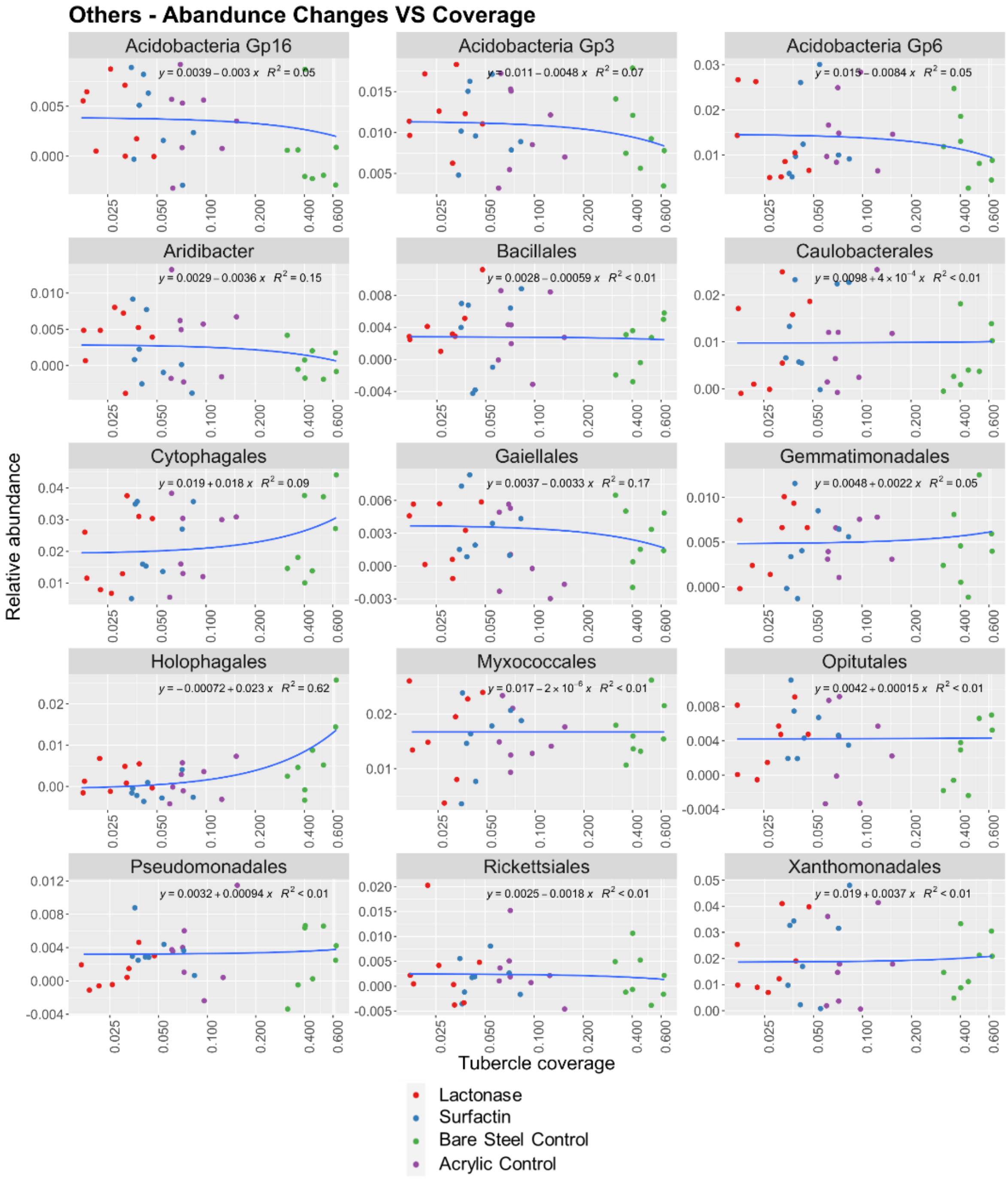

**Fig S15**. Log scale plots showing the correlation between the relative abundance of bacterial orders and the tubercle coverage for the 40 most abundant bacterial orders by total sequence counts for treatment and control samples. The bacterial orders are grouped by IOB, SRB and others. All sites and depths are combined in the analysis.

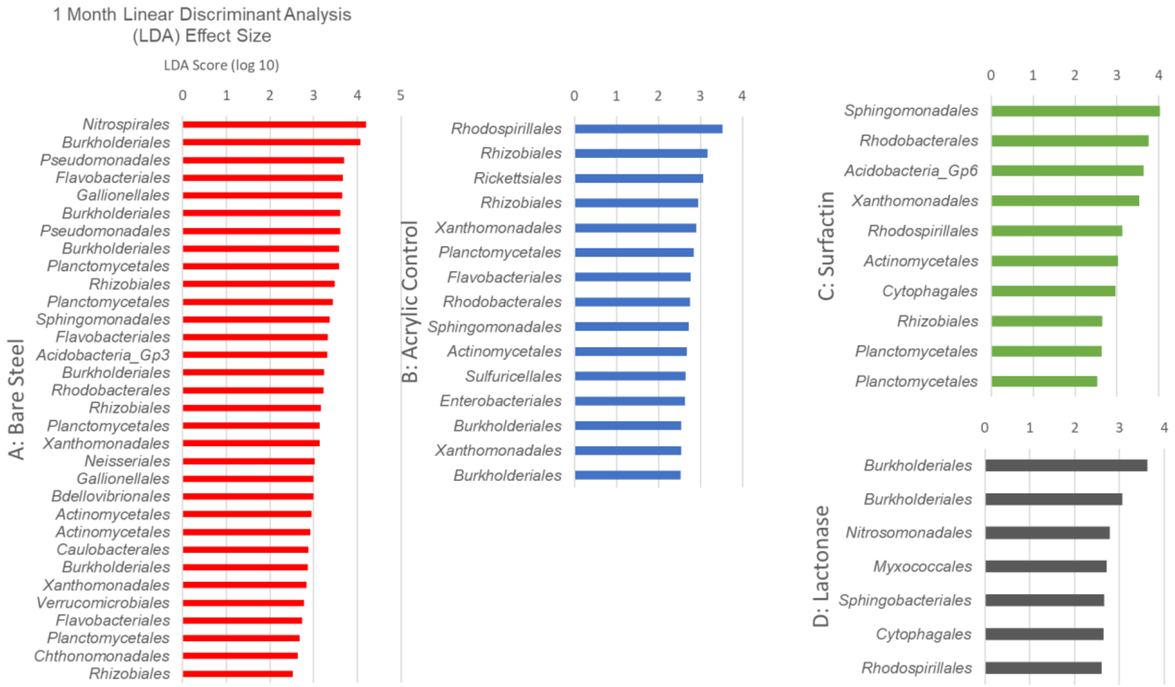
1 Month Linear Discriminant Analysis (LDA) Effect Size
LDA Score (log 10)
A: Bare Steel
Nitrospirales
Burkholderiales
Pseudomonadales
Flavobacteriales
Gallionellales
Burkholderiales
Pseudomonadales
Burkholderiales
Planctomycetales
Rhizobiales
Planctomycetales
Sphingomonadales
Flavobacteriales
Acidobacteria_Gp3
Burkholderiales
Rhodobacterales
Rhizobiales
Planctomycetales
Xanthomonadales
Neisseriales
Gallionellales
Bdellovibrionales
Actinomycetales
Actinomycetales
Caulobacterales
Burkholderiales
Xanthomonadales
Verrucomicrobiales
Flavobacteriales
Planctomycetales
Chthonomonadales
Rhizobiales
B: Acrylic Control
Rhodospirillales
Rhizobiales
Rickettsiales
Rhizobiales
Xanthomonadales
Planctomycetales
Flavobacteriales
Rhodobacterales
Sphingomonadales
Actinomycetales
Sulfuricellales
Enterobacteriales
Burkholderiales
Xanthomonadales
Burkholderiales
C: Surfactin
Sphingomonadales
Rhodobacterales
Acidobacteria_Gp6
Xanthomonadales
Rhodospirillales
Actinomycetales
Cytophagales
Rhizobiales
Planctomycetales
Planctomycetales
D: Lactonase
Burkholderiales
Burkholderiales
Nitrosomonadales
Myxococcales
Sphingobacteriales
Cytophagales
Rhodospirillales

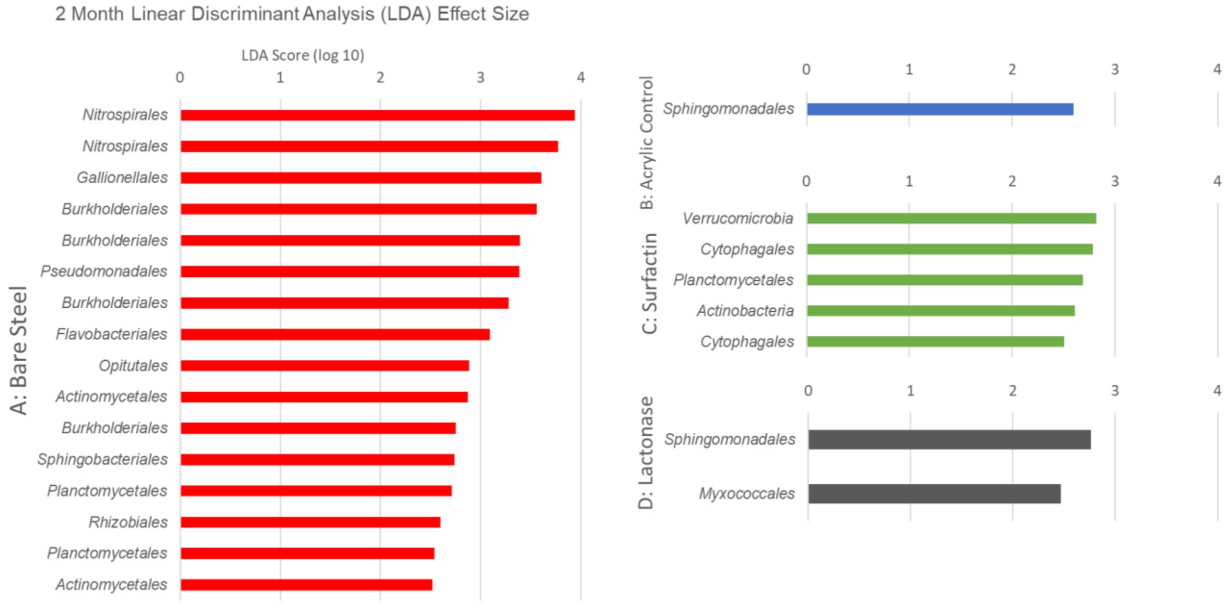
2 Month Linear Discriminant Analysis (LDA) Effect Size
LDA Score (log 10)
A: Bare Steel
Nitrospirales
Nitrospirales
Gallionellales
Burkholderiales
Burkholderiales
Pseudomonadales
Burkholderiales
Flavobacteriales
Opitutales
Actinomycetales
Burkholderiales
Sphingobacteriales
Planctomycetales
Rhizobiales
Planctomycetales
Actinomycetales
B: Acrylic Control
Sphingomonadales
C: Surfactin
Verrucomicrobia
Cytophagales
Planctomycetales
Actinobacteria
Cytophagales
D: Lactonase
Sphingomonadales
Myxococcales

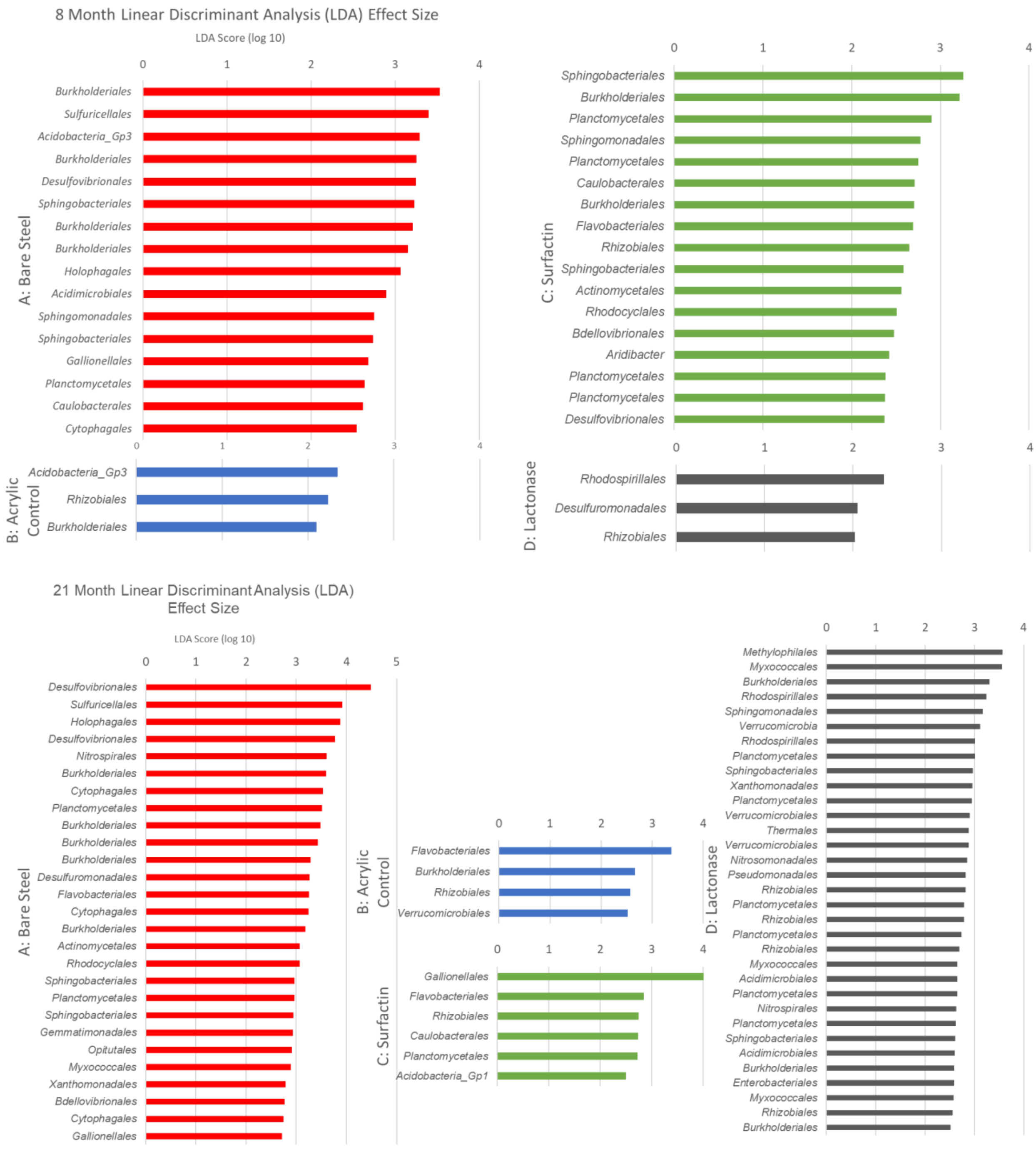

**Fig S16**. The linear discriminant analysis (LDA) effect size (LEfSe) analysis identified bacterial orders that responded significantly to the different treatments during the four sampling periods. Each bar represents the effect size of an OTU. All sites and depths were combined in this analysis. Relative abundance was significant when $P < 0.05$, logarithmic LDA score $\geq 2$.
